# Temporal Microbiome Changes in Axolotl Limb Regeneration: Stage-Specific Restructuring of Bacterial and Fungal Communities with a *Flavobacterium* Bloom During Blastema Proliferation

**DOI:** 10.1101/2024.03.07.583834

**Authors:** Hanne Altın, Büşra Delice, Berna Yıldırım, Turan Demircan, Süleyman Yıldırım

## Abstract

The intricate relationship between regeneration and microbiota has recently gained attention, spanning diverse model organisms. Axolotl (*Ambystoma mexicanum*) is a critically endangered salamander species and a model organism for regenerative and developmental biology. Despite its significance, a noticeable gap exists in understanding the interplay between axolotl regeneration and its microbiome. Here, we analyze in depth bacterial 16S rRNA amplicon dataset that we reported before as data resource and profile fungal community by sequencing ITS amplicons at the critical stages of limb regeneration (0-1-4-7-30-60 days post amputation, “dpa”). Results reveal a decline in richness and evenness in the course of limb regeneration, with bacterial community richness recovering beyond 30 dpa unlike fungi community. Beta diversity analysis reveals precise restructuring of the bacterial community along the three phases of limb regeneration, contrasting with less congruent changes in the fungal community. Temporal dynamics of the bacterial community highlight prevalent anaerobic bacteria in initiation phase and *Flavobacterium* bloom in the early phase correlating with limb blastema proliferation. Predicted functional analysis mirrors these shifts, emphasizing a transition from amino acid metabolism to lipid metabolism control. Fungal communities shift from *Blastomycota* to *Ascomycota* dominance in the late regeneration stage. Our findings provide ecologically relevant insights into stage specific role of microbiome contributions to axolotl limb regeneration.

## 1 | INTRODUCTION

The axolotl (*Ambystoma mexicanum*), a species facing critical endangerment in its natural habitat, is a member of the *Ambystoma* clade within the mole salamander family (*Ambystomatidae*). Distinguishing salamanders from other amphibians is their remarkable capacity for complete regeneration, encompassing not only limbs and appendages but also certain internal organs. The intricate cellular and molecular mechanisms involved in the extensively researched limb regeneration of axolotls make it a topic of significant scientific interest ^1, 2^.

To regenerate its limbs, the axolotl employs a fascinating process involving the generation of blastema cell mass, resembling the embryonic limb bud. This intricate procedure is significantly influenced by innervation, where the blastema acts as the driving force for redevelopment through a combination of growth and redifferentiation. Despite its complexity, limb regeneration in axolotls unfolds systematically in a stepwise manner. Indeed, it is well established that the regeneration progresses over distinct interdependent stages according to morphogenesis ^3^. These sequential phases encompass wound healing, dedifferentiation (highlighted by blastema establishment) and re-development stages of regeneration, respectively ^4–6^. The cell lineage- and regeneration stage-specific expression patterns of various morphogenesis genes are critical for successful limb regeneration and the early steps are highly critical for determining the extent of regenerative response after limb amputation. More specifically, after limb amputation, a specialized wound epidermis forms within ~24 hours through cell migration. Wound epidermis is structurally and molecularly distinct from fully differentiated, intact epidermis. Innervated wound epidermal cells then thicken the tissue, dermal cells migrate beneath it, and proliferates. In the days following re-epithelialization, progenitor cells enter into the cell cycle as well as the accumulate at the tip of the stump, beneath the wound epidermis. These progenitor cells may originate from stem cells or by dedifferentiation. Epigenetic regulation controls both wound epidermis formation without scarring and epithelial-to-mesenchymal transition ^1, 7, 8^. Together, the activated cells accumulated at the tip of the stump give rise to highly proliferative blastema cells ^1, 7, 9^.

Timing of these events may slightly vary in different salamander species but in axolotl the movement of cells from the dermis into the early limb blastema begin to migrate beneath the wound epithelia for about 5 days post amputation (dpa0-5: “initiation phase”). Next, the wound epithelium thickens and forms an apical epithelial cap (AEC), which promotes the generation of a population of undifferentiated mesenchymal cells called a ‘blastema’, which is formed through a ‘dedifferentiation’ process from differentiated tissues (dpa6-20: “early phase”). In the re-development stage, the blastema continues to grow distally via cellular proliferation until the completion of regeneration (dpa21-30: “late phase”) ^4–6, 10, 11^.

Animal microbiomes play key roles in an animal’s development, health, behavior, and protection from pathogens ^12–15^. Recently, the impact of the host microbiome on regenerative ability of its tissues, organs or even whole-body has gained significant attention even though this is still relatively a new area of research in the field of microbiomes. Indeed, several species, renowned for their regenerative capabilities, such as planarians, sea cucumber, and zebrafish, have been employed in research to investigate whether the microbiome plays a role in regulating the regenerative potential of these hosts or is directly engaged in the regeneration process. These studies revealed that some bacterial species were shown to be causally linked with the regenerative ability of the model animals. For instance, *Pseudomonas* and *Aquitalea* species on Planaria, *Erwinia carotovara* on fruit fly, *Aeromonas veronii* on Zebrafish, *Akkermansia muciniphila* and some probiotic *Lactobacillus species* in intestinal cell proliferation ^16^. The microbiome, therefore, influence tissue regeneration by modulating immune responses and inflammation, and potentially contributing to the activation of regenerative pathways.

The axolotl is a permanently aquatic animal and its amputated limb regenerates against the backdrop of microbial communities of the aquaria. This raises the question as to whether and how axolotl’s gut and skin microbiota and water microbiota in the aquarium tank impacts the limb regeneration. Remarkably, however, reports on the impact of microbiome on this highly regenerative animal, the axolotl, and salamanders at large, is very limited, leaving significant knowledge gap. We previously generated high quality 16S rRNA dataset from a longitudinal sampling of axolotl limb tissues over the course of regeneration stages and exceptionally presented the dataset to the interest of the axolotl scientific community ^17^.

In this study, we comprehensively analyze the diversity and temporal dynamics of both bacterial and fungal communities within the same experimental setup. We hypothesized that regenerating limb microbial communities are distinct from the environment; host factors create unfavorable environment for the colonization of pathogens and induce the microbial communities to progressively adapt to evolving micro niches in the course of axolotl limb regeneration. Our results demonstrate that both bacteria and fungi communities dramatically restructure over the course of axolotl limb regeneration stages and show lack of resilience at the fully regenerated stage. Our findings also provide evidence that the host limb tissues select microbial taxa that can colonize in the regenerated tissues.

## 2 | MATERIAL AND METHOD

### 2.1 | Ethical Statement and Experimental Design

The ethical statement and experimental design were previously described in our previous report^17^. Briefly, this study was conducted at the Regenerative and Restorative Medicine Research Center (REMER) of the Istanbul Medipol University. Experimental protocols and animal care conditions were approved by the local ethics committee of the Istanbul Medipol University (the authorization number 38828770-604.01.01-E.10834). The founders of the initial axolotls colony were procured from the University of Kentucky, USA and reared in optimized conditions^18^ in aquaria in animal care facility of the university.

A total of 54 axolotls were selected at random from a group of siblings. The randomly selected animals were then individually housed in aquariums filled with Holtfreter’s solution at a constant temperature of 18 ± 2 °C. They were fed with standard food (JBL Novo LotlM) once daily. The experimental design, shown in Figure 1, included nine animals per group, divided into three biological replicates (R1, R2, R3) to assess reproducibility. During the study, each animal was kept in a separate aquarium. Limb amputation involved anesthetizing the axolotls with 0.1% Tricaine methane sulfonate (Cat. No. E10521 or MS-222, Sigma-Aldrich, St. Louis, MO, USA) and amputating the right forelimb at the mid-zeugopod level.

### 2.2. | Sample Collection

After amputation, animals were randomly assigned to six groups, simulating axolotl limb regeneration phases: initiation (dpa 0 and dpa 1), early (dpa 4 and dpa 7), and late (dpa 30 and dpa 60). Tissue samples from three animals were pooled to reduce variation, cryopreserved in liquid nitrogen, and stored at −80 °C until genomic DNA isolation. Sampling specifics included 1-mm tissue around the cut site for dpa 0 and dpa 1, removal of newly formed blastema and 0.5 mm posterior tissue for dpa 4 and dpa 7, and collection around the original cut site for dpa 30 and dpa 60 to analyze microbiota in regenerated tissues.

To explore microbiota colonization in axolotl limbs originated from Holtfreter’s solution, we obtained 100 ml water samples from 9 separate aquaria throughout the experiment. We combined samples from the beginning (day0-R1), middle (day30-R2), and end (day60-R3) of the experimental timeline, forming the “aqua” control group with three replicates (R1, R2, R3; see Fig. 1a).

### 2.3. | DNA Extraction and Amplicon Sequencing

Genomic DNA was isolated from skin tissue samples using the DNeasy Blood and Tissue Kit (Qiagen, Catalog No. 69504), while the Metagenomic DNA Isolation Kit (Epicentre, catalog no: MGD08420) was employed for Aquarium water samples. The concentration of the isolated genomic DNA was measured using the Qubit 2.0 Fluorometer (Thermo Fisher Scientific, MA, USA). The internal transcribed spacer 1 (ITS1) region primers (ITS1-30F: GTCCCTGCCCTTTGTACACA and ITS1-217R: TTTCGCTGCGTTCTTCATCG) were selected from a previously published report ^19^ and V3-V4 region of 16S rRNA gene was amplified in the previous study as described before^17^. To ensure DNA integrity, an aliquot of the samples was run on a 1.0% agarose gel. For each sample, PCR was performed in a total volume of 25 μl with two replicates, including approximately 12.5 ng of purified DNA template and 2x KAPA HiFi HotStart Ready Mix. The PCR conditions consisted of an initial denaturation at 95 °C for 3 minutes, followed by 25 cycles of denaturation at 95 °C for 30 seconds, annealing at 55 °C for 30 seconds, and extension at 72 °C for 30 seconds. The final extension was at 72 °C for 5 minutes. Negative controls in genomic DNA isolation and PCR steps were included to control potential contamination and sequenced along with the other samples.

Amplicons were purified using the Agencourt AMPure XP purification system (Beckman-Coulter Cat. No. A63881, USA). A second PCR, with 8 cycles, was conducted using sample-specific barcodes, and the obtained amplicons were purified again. Equimolar concentrations from each library were pooled and sequenced on an Illumina MiSeq sequencer using the MiSeq Reagent Kit v2 (500 cycles). Raw data from this sequencing effort can be found in the NCBI Sequence Read Archive (Accession number: PRJNA1073213).

### 2.4. | Sequence Processing Pipeline and Taxonomic Classification

Pair-end reads of the amplicons were first merged using cutadapt^20^ after removing primers and low quality sequences. The merged clean sequence reads were then uploaded to the Nephele platform (v.2.30, http://nephele.niaid.nih.gov) to run DADA-ITS pipeline and 16S rRNA pipeline on default parameters and their associated denoising and removing chimeric reads. For taxonomic assignment of the reads the SILVA small-subunit rRNA sequence database (v.132)^21^ for bacteria and the database for Fungi^22^ were used. To run the Phylogenetic investigation of communities by reconstruction of unobserved states (PICRUSt2)^23^ analysis we locally reran QIIME2 on Greengenes taxonomy. The resulting amplicon sequence variants (ASVs) table was filtered to exclude chloroplasts, mitochondria, archaea, eukaryotes, or unknown reads were eliminated. As the number of sequences per sample were quite imbalanced, we rarefied ASV tables before starting the downstream analyses and applied center log ratio (CLR) transformation. Downstream analysis of ASVs was carried out using the package Phyloseq^24^ in R version 4.1.0.

### 2.5. | Community Structures and Diversity

The alpha diversity at each sampling time point was calculated using Chao1, Shannon, and Simpson metrics. The Kruskal-Wallis (KW) test and posthoc Dunn’s test were used to identify alpha diversity differences between samples. To test whether bacterial and fungal communities differ in community composition and structure we employed Canonical analysis of principal coordinates (CAP) based on Bray-Curtis distance matrix. Statistically significant differences among life stages were determined with permutational ANOVA (PERMANOVA) performed with adonis function. To test pairwise group differences were tested using pairwise.adonis wrapper function for multiple testing 999 in vegan R package and p values were corrected with False Discovery Rate (FDR) for multiple testing. Indicator species analysis was performed for each in axolotl limb regeneration samples using the indicspecies package in R. Spearman rank-based correlation coefficients were calculated using from the Hmisc R package. Moreover, pheatmap, lattice, UpSetR and ggplot2 R packages were used for visualizations. Longitudinal changes of taxa of interest was plotted using LOESS() function that fits local polynomial regression within stats (v3.6.2) and ggplot2 R packages.

## 3 | RESULTS

In our previous work, we provided a comprehensive overview of the experimental design employed in this study, as well as the 16S rRNA dataset generated and shared with the scientific community for further analysis or meta-analysis^17^. Expanding on this groundwork, we broadened our inquiry by analyzing ITS1 amplicons from identical archival gDNA samples to characterize the temporal changes of the mycobiome during the various stages of limb regeneration. These stages were categorized as dpa0-dpa1 for the initiation phase, dpa4-dpa7 for the early phase, and dpa30-dpa60 for the late phase. In this study, we provide a comprehensive analysis of both bacterial and fungal communities simultaneously.

Briefly, we meticulously monitored the amputated axolotls for 60 days and longitudinally collected a total of n=21 samples (comprising 18 limb tissue samples from n=54 adult axolotls and 9 individual aquaria samples with replicates) in total (Figure 1). Subsequent to preprocessing ITS amplicon sequences, conducting quality checks, and removing chimeric sequences using the DADA2 pipeline we obtained 238 amplicon sequence variants (ASVs) with sufficient sequencing depth (28978 reads/sample in the rarefied (ASVs) table). The bacteria ASV table included 116220 reads/sample in the rarefied ASV table.

### 3.1. | Dynamic restructuring of bacterial and fungal diversity across regeneration stages

We compared species richness and diversity using a variety of alpha-diversity indices including Chao1, Shannon, and Simpson and observed consistent decreasing trend in all metrics of both bacterial diversity (Figure 2A) and fungal diversity (Figure 2B). Unlike fungal community bacterial richness did not reach significance level among samples despite the apparently decreasing trend as measured by Observed species and Chao1 indices (KW, p>0.05, Table S1.1). In contrast, Shannon and Simpson indices steeply increased at dpa30 and dpa60 (late phase or re-development of the limbs, KW, P=0.01375 for Shannon and P=0.01423 for Simpson), suggesting bacteria community colonizing the re-growing limb skin was more evenly restructured than fungal community despite the decrease in the number of species. Indeed, Dunn’s posthoc pairwise comparisons for each alpha diversity index for bacterial community showed significant differences between all sample types except between the samples of the initiation phase and late phase while this pattern was absent for the fungal community samples (Table S2.1-S2.2).

Next, we analyzed temporal dynamics of both bacteria and fungi communities using Bray Curtis distance and Canonical Analysis of Principal Coordinates (CAP) (Figure 3A), a constrained ordination that maximizes the differences among a priori groups. We reproduced our prior findings^17^, indicating distinct separation among bacterial communities at each sampling time point (PERMANOVA, P=0.001, pseudo-F=7.191), a result further bolstered by statistically significant hierarchical clustering (see Figure 3B) and confirmed by the SIMPROF significance test (significance level 5%, 999 permutations). These results were mirrored by the CAP and cluster analysis predicted metagenome functional pathway abundances (PICRUSt2) (Figure 3C-D, PERMANOVA, pseudo-F=10.461, P=0.001 and SIMPROF (5%, 999 permutations, respectively). Interestingly, despite the significant differences in distances between centroids of fungal community samples (Pseudo-F=2.715, P=0.004, see Figure 4A), both hierarchical clustering (Figure 4B) and pairwise PERMANOVA analysis (using Monte Carlo permutation) revealed that fungal communities did not exhibit significant separation at any pairs of dpa timing points (p>0.05), unlike the pairwise contrasts observed in bacterial communities (p<0.05) (Table S3). This analysis suggests that the spatial and temporal dynamics of the fungal community in response to regeneration are distinct from those of the bacterial community.

### 3.2. | Regeneration stages prompt restructuring of bacterial communities, triggering *Flavobacterium* bloom

The heatmap of Top20 most abundant bacteria taxa at genus level showed conspicuous clustering of co-abundant taxa by the limb regeneration stages of axolotl (Figure 5). In the initiation stage (dpa0-1) largely anaerobic gut bacteria (*Bacteroides, Akkermansia, Paenarthrobacter, Odoribacter, Parabacteroides,* and unclassified species within the family of *Ruminococcoaceae* and *Lachnospiraceae*) were clustered together while in the early phase (dpa4-7) a bloom of *Flavobacterium* was observed. Finally, in the late phase (dpa30-60) the bacterial profile continued to evolve, giving rise to the potentially opportunistic bacteria (e.g. *Staphylococcus, Pseudomonas)* or taxonomically closely related species that are often isolated from soil freshwater creeks, lakes (e.g. *Elizabethkingia* and *Chryseobacterium* in the order of *Flavobacteriales*) that colonize on the skin of the re-growing limbs.

Furthermore, we employed LOESS regression, which uses weighted local polynomials to model species abundance data across time, to demonstrate longitudinal shifts of Top20 taxa. Clearly, in the early phase (dpa4-7) a bloom of *Flavobacterium* persisted during the critical early stage that allows for formation and growth of blastema (Figure 6A). A significant increase in the abundance of *Chryseobacterium* over dpa60 was also remarkable. Additionally, we performed indicator species analysis to unravel indicator taxa at each sampling time point (Figure 6B), which further supported that *Flavobacterium* is indeed an indicator taxon of the early phase, a critical window for blastema to start proliferating. Notably, *Aquabacterium* and *Acinetobacter* were also found to be an indicator taxa within the early phase although their abundances were low. To answer the question whether functional pathway abundances of bacteria communities follow similar trend as taxa in the course of limb regeneration stages we plotted the prediction of metagenome functional pathway abundances (PICRUSt2) in a heatmap (Figure 7). Remarkably, we observed clustering of pathways by limb regeneration stages, mirroring the taxa abundance trend. The pathways in the initiation phase were found to be largely associated with amino acid and nucleic acid biosynthesis while lipid biosynthetic pathways dominated the following early phase. The late phase did not show clustered pathways. Correlation analysis between abundances of bacteria taxa and pathways highlighted *Chryseobacterium* and *Flavobacterium* highly significantly associated with the amino acid and lipid pathways, respectively (r=0.40, p<0.05, Figure S1). Together, the diversity and co-abundance of bacteria communities and their clustered pathways point toward stage specific restructuring of bacterial communities across the limb regeneration while restructuring of the fungal communities does not closely follow the regeneration stages.

### 3.3. | Unexpected complexity of fungal communities during the course of limb regeneration

The lack of obvious restructuring trend within fungal community was also manifested in the taxonomic diversity of fungal communities. In spite of significant shift in community composition from Basidiomycota dominance to Ascomycota prevalence (Figure 6A) genus level fungal taxa did not cluster by the stages of limb regeneration (Heatmap, Figure 6B). These findings prompted us to investigate degree of temporal relationship between bacterial and fungal communities. We therefore performed Procrustes analysis and Mantel correlation between the Bray-Curtis distance matrices for both communities. Interestingly, both Procrustes and Mantel analyses identified a moderately significant relationship between the two datasets (Procrustes correlation=0.73, *p*=0.001 and Mantel test r=0.69, *p*=0.001), suggesting both communities influence each other during regeneration. In addition, we found significant correlations (r=±0.40, P*<0.05)* between bacteria and fungi taxa (Figure 7). In addition to *Flavobacterium, Unclassified Neisseriaceae, Chryseobacterium, Elizabethkingia, Unclassified Burkholderiaceae*, and *Akkermansia* exhibited significant correlations with fungal taxa. Notably, *Flavobacterium* and *Unclassified Burkholderiaceae* showed predominantly negative correlations. Among fungal taxa, *Talaromyces, Aspergillus,* and *Stachybotrys* displayed significant correlations with bacterial taxa.

Finally, since Aqua samples were placed farther away from the limb tissue samples in CAP figures we compared the number of unique and shared taxa between limb tissue and aqua samples (Figure 8A-B). The upset figures shows both bacterial and fungal communities are substantially unique at each sampling points (dpa0-dpa60) and few taxa persisted as shared species between these time points, pointing towards highly dynamic temporal shifts of the community compositions over the course of the limb regeneration. As shown in Figure 8A the highest number of unique bacterial species assembled at dpa0 (562 ASVs) and dpa30 (535 ASVs) and 36 ASVs are common among the sampling points. As expected, only 14 ASVs are present both in Aqua water and all limb tissues, suggesting the temporal dynamics of the microbiota colonizing regenerating tissues is due largely to the host selection although microbe-microbe interactions could also contribute to unique community profiles.

## 4 | DISCUSSION

The axolotl, with its remarkable ability to regenerate limbs and organs due to sustained neoteny, has strongly attracted the attention of scientific community for translational medicine research^3^. Despite its recently sequenced expansive genome^25^, studies on the axolotl microbiome are very limited. In this pioneering study, we unveil, for the first time, the substantial restructuring of both fungal and bacterial communities throughout the intricate process of limb regeneration. Our findings illuminate the emergence of temporally unique community compositions at sampling days that correspond to the widely accepted three phases of limb regeneration in axolotls. Remarkably, as the axolotl’s limbs undergo full regeneration, the microbial communities inhabiting the limb skin undergo significant turnover. This suggests that the axolotl limb skin microbiome lacks resilience in response to the regenerative process. Such insights into the dynamic interplay between microbial communities and the regenerative capacity of axolotls shed new light on the intricate mechanisms underlying limb regeneration. Moreover, they offer valuable implications for translational research aimed at harnessing the regenerative potential of axolotls for therapeutic purposes in humans.

One of the most striking observations on limb microbiota is regeneration-induced changes in richness and evenness of limb microbiota as evidenced by significantly decreased alpha diversity metrics for both bacteria and fungi communities. Notably, Shannon and Simpson indices calculated for bacteria communities reversed the downward trend at dpa30 unlike fungal communities. As the axolotl regrows its regenerated limb during this stage, it becomes apparent that the fungal community struggles to recover its original diversity post-treatment. Notably, however,

Fungi and bacteria often have different life cycles and growth rates, and regeneration stages might favor bacterial growth or alters resource availability in a way that benefits bacteria over fungi, contributing to the disparity in alpha diversity metrics. Moreover, although fungi community significantly shifts during the course of limb regeneration as manifested by the PERMANOVA main test the specific pairwise differences between individual groups were not statistically significant. Even though this finding can be attributed to the small sample size (3 replicates) for each pairwise comparison leading to limited power to detect true effects the hierarchical clustering of the samples was not congruent with the stage specific separation. Axolotl limb fungi may therefore have yet unknown subtle ecological complexity in responding to traumatic amputation and regeneration processes.

*Flavobacterium*, among other microbiota, warrants special attention during the course of limb regeneration as our results indicated this bacterium blooms across dpa4-dpa7 where the wound epithelium thickens and forms an apical epithelial cap, which promotes the generation of a population of undifferentiated mesenchymal cells called a ‘blastema’. As a result, signals from the host and environmental cues can potentially promote or hinder differentiation and/or proliferation of the blastema cells in this critical stage. *Flavobacterium* belongs to the family *Flavobacteriaceae* in the phylum *Bacteroidota.* It is a Gram-negative genus comprising 270 species, ubiquities in the environment, including soil, seawater, freshwater, plants, and glaciers ^26, 27^. Many species of the genus *Flavobacterium* are capable of plant growth-promoting ability, hydrolyzing organic polymers, such as complex polysaccharides, cold-adapted species produce polyunsaturated fatty acids ^26^, although some species (such as *F. columnare* and *F. psychrophilum*) reportedly cause infectious diseases in freshwater fish^28, 29^. Nonetheless, considering axolotl was able to complete limb regeneration in this experiment in spite of the bloom of this bacterium the unknown species of *Flavobacterium* in this context may be positively contributing to the regeneration. While its putative contribution to regenerative mechanisms and epigenetic and antigenic interactions with blastema cells cannot be ruled out evidence in the published literature support that beneficial cutaneous bacteria on amphibians protect against the lethal disease chytridiomycosis caused by *Batrachochytrium dendrobatidis* (Bd) and salamanders ^30–34^. In this study our analysis of fungal community did not detect Bd with the caveat that the majority of the ITS reads were not assigned to Unclassified Fungi. We therefore reasoned that either ITS primers did not amplify Bd, if present, or *Flavobacterium* might compete against unknown fungal species that could potentially impede regeneration mechanisms. Moreover, the prevalence of *Flavobacterium* exhibited a robust correlation with pathways related to lipid metabolism (see Figure S1), indicating a potential role for this bacterium in influencing regeneration through lipid metabolic processes. Interestingly, a recent study analyzed metabolomic profile of axolotl blastema at dpa11^35^. Lipid metabolites such as sphingomyelin and sphingosine, along with polyamine and pyrimidine metabolism, were found to be enriched in blastema tissue. A recent cutaneous microbiota study on Ozark hellbender salamander subspecies (*Cryptobranchus alleganiensis bishopi*), suffering from chronic wounds within their aquatic habitat has uncovered interesting findings^36^. The study showed that *Acidovorax sp., Flavobacterium sp., Aquabacterium sp., Chryseobacterium sp.,* and *Acinetobacter sp.* were found to have higher relative abundance within these wounds compared to intact skin. These results are consistent with our own research, as all four species were identified as indicator species during axolotl limb regeneration studies. Nonetheless, further investigation is needed to determine whether these bacterial species, particularly *Flavobacterium*, are associated with regenerative capacity of axolotl limb.

The stage specific restructuring of bacteria and fungi communities across the limb regenerating phases is fascinatingly chaotic and unstable as we found largely unique ASVs at almost every dpa timepoint (Figure 8A-B). Given that wound epidermis covers the wound within a few hours of amputation and Apical Epithelial Cap (AEC) cells quickly develops in the initiation phase the newly colonizing microbial communities at each dpa point would first have a chance to briefly interact with the wound epidermis but subsequently AEC cells during the critical stage of blastema proliferation. Even though the chaotic stage specific restructuring of microbial communities can be due partly to microbial resource competition the host factors is likely in control of the community composition, provides the microbes less than favorable environment to form continuous biofilms at least until the re-developmental stage is completed. The host factors include the proinflammatory immune response to amputation and regeneration. Indeed, macrophages infiltrate amputation site about the same time wound epidermis covers the amputation site and depletion of macrophages impedes limb regeneration ^37^. Theoretically, genetic and epigenetic signaling between macrophages, AEC cells, nerve cells, blastema, oxidative-stress, and ultimately redecoration of limb skin with protective mucus layer all would have impact restructuring of microbial community attempting to re-colonize regenerating tissues.

## Conclusions

Our findings offer compelling evidence for the first time that both bacterial and fungal communities undergo restructuring across the widely accepted three major phases in axolotl limb regeneration. However, it is noteworthy that the restructuring of the fungal community is less precisely congruent with the timing of these major phases. The notable bloom of *Flavobacterium* during the critical stage of blastema proliferation, along with the compositional shift of fungi from Basidiomycota dominance in the early phase to Ascomycota dominance in the late phase, raises intriguing questions about their potential impact on regenerative mechanisms in the limb tissues through unknown mechanisms. Future research, therefore, should investigate the contribution of microbial communities to regenerating tissues, specifically the cross talk between *Flavobacterium* and macrophages, and AEC cells. By unraveling these intricate interactions, we can gain deeper insights into the role of microbiota in the complex process of tissue regeneration, paving the way for potential therapeutic interventions that harness the theraputic potential of microbial communities in enhancing tissue repair and regeneration.

## AUTHOR CONTRIBUTIONS

Conceptualization: SY and TD. Data curation: HA and SY. Formal analysis: SY, BD, and HA. Methodology and resources: TD, BY, and SY. Writing – original draft: SY, Writing – review & editing: SY, TD, HA, and BY. All authors read and approved the final manuscript.

## ACKNOWLEDGMENTS

The financial support for this study was provided by the Medipol University Research Fund. We extend our gratitude to Ms. Ayça Önal and Ms. Damla Yay for their invaluable assistance in capturing axolotl limb photographs.

## DATA AVAILABILITY STATEMENT

The data that support the findings of this study are openly available in NCBI Bio project at https://www.ncbi.nlm.nih.gov/bioproject/….., reference number …..

---

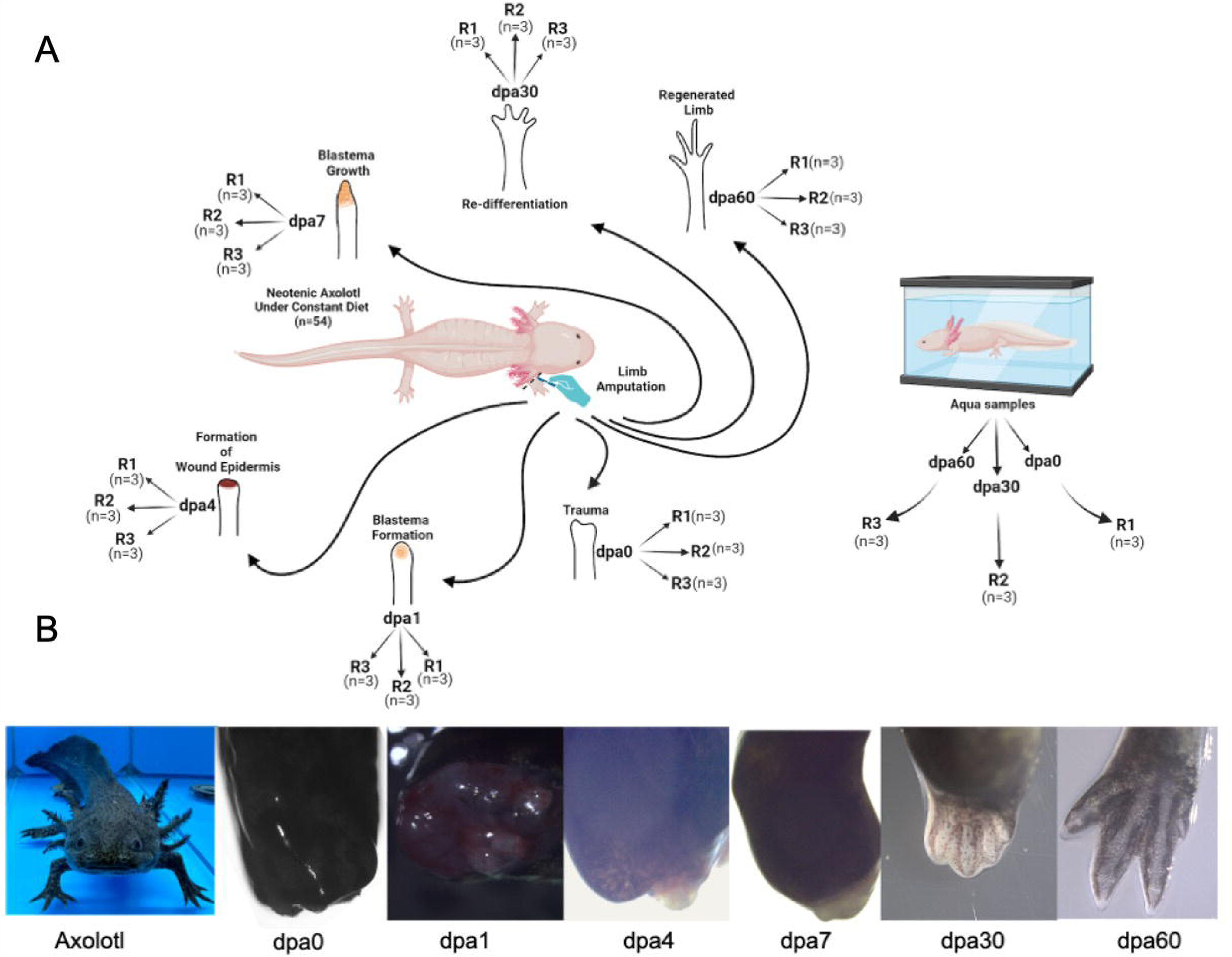

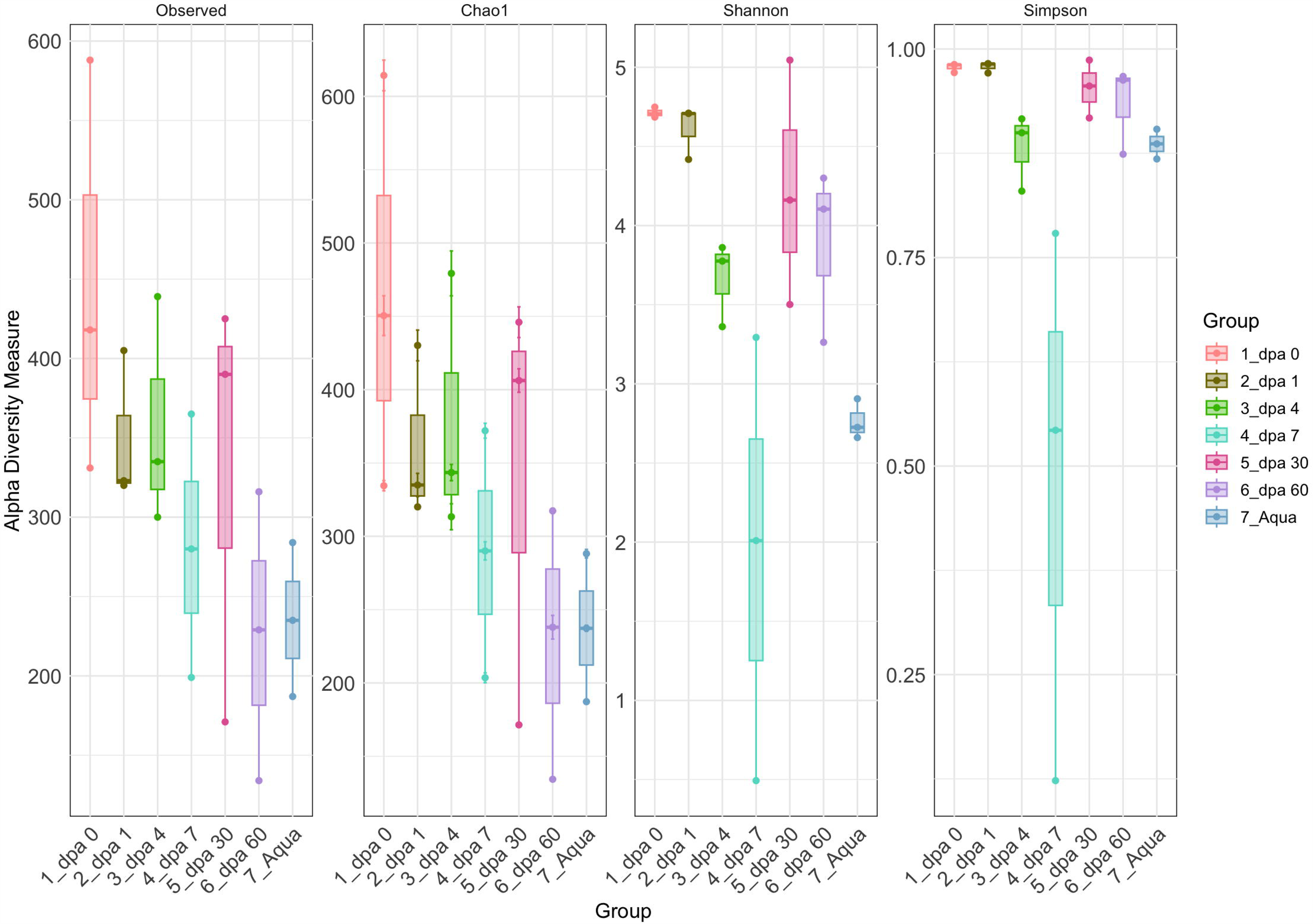

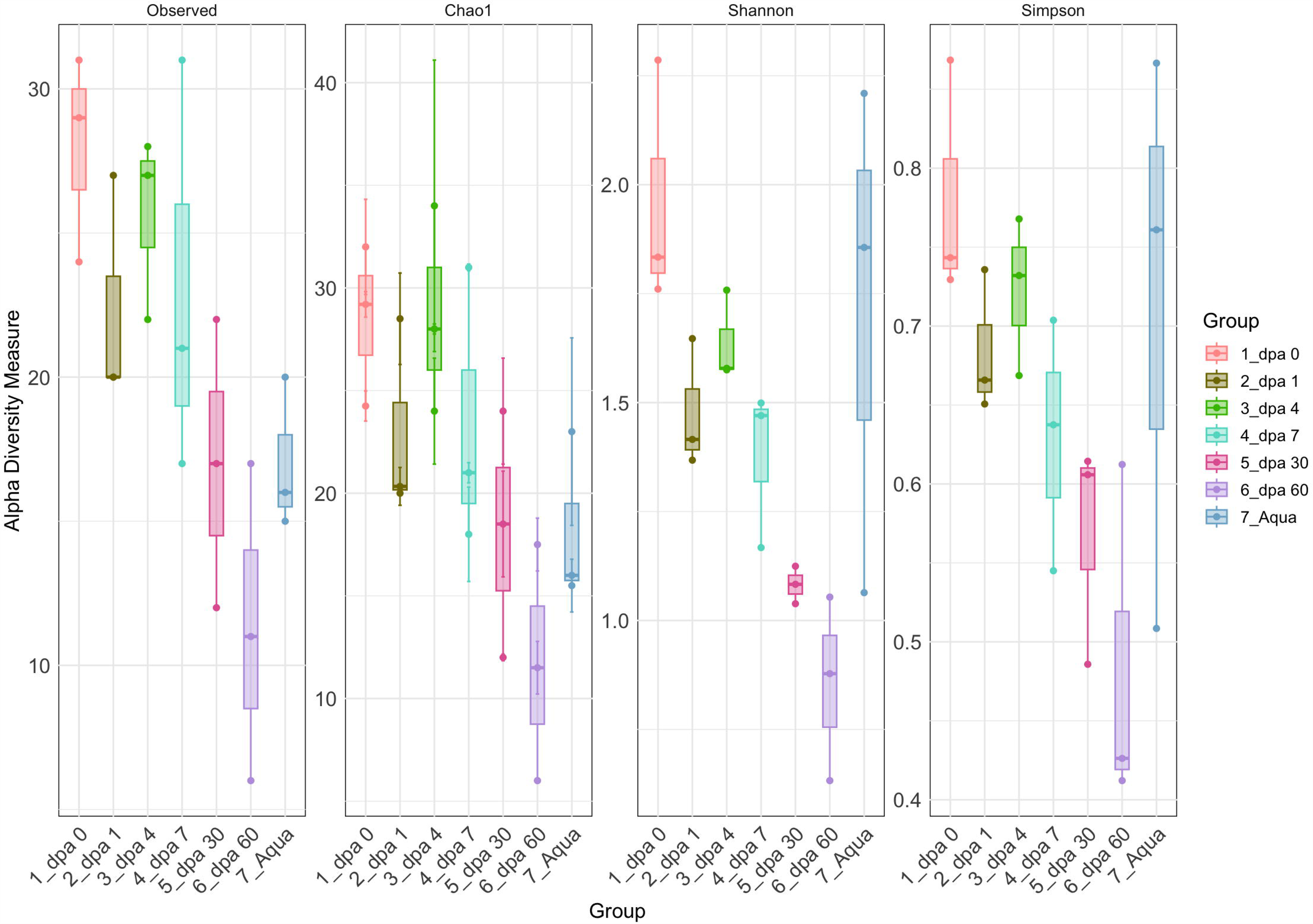

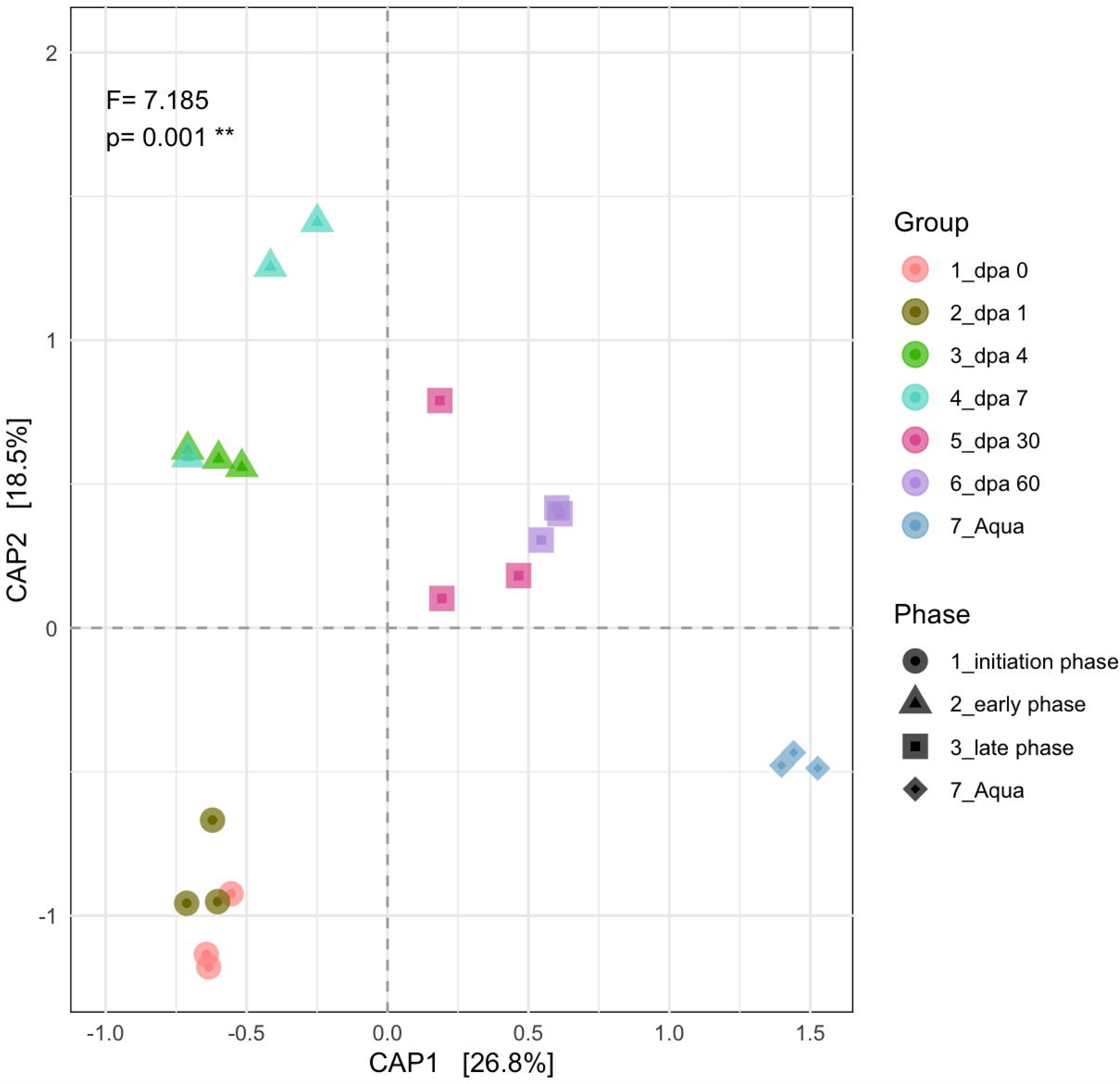

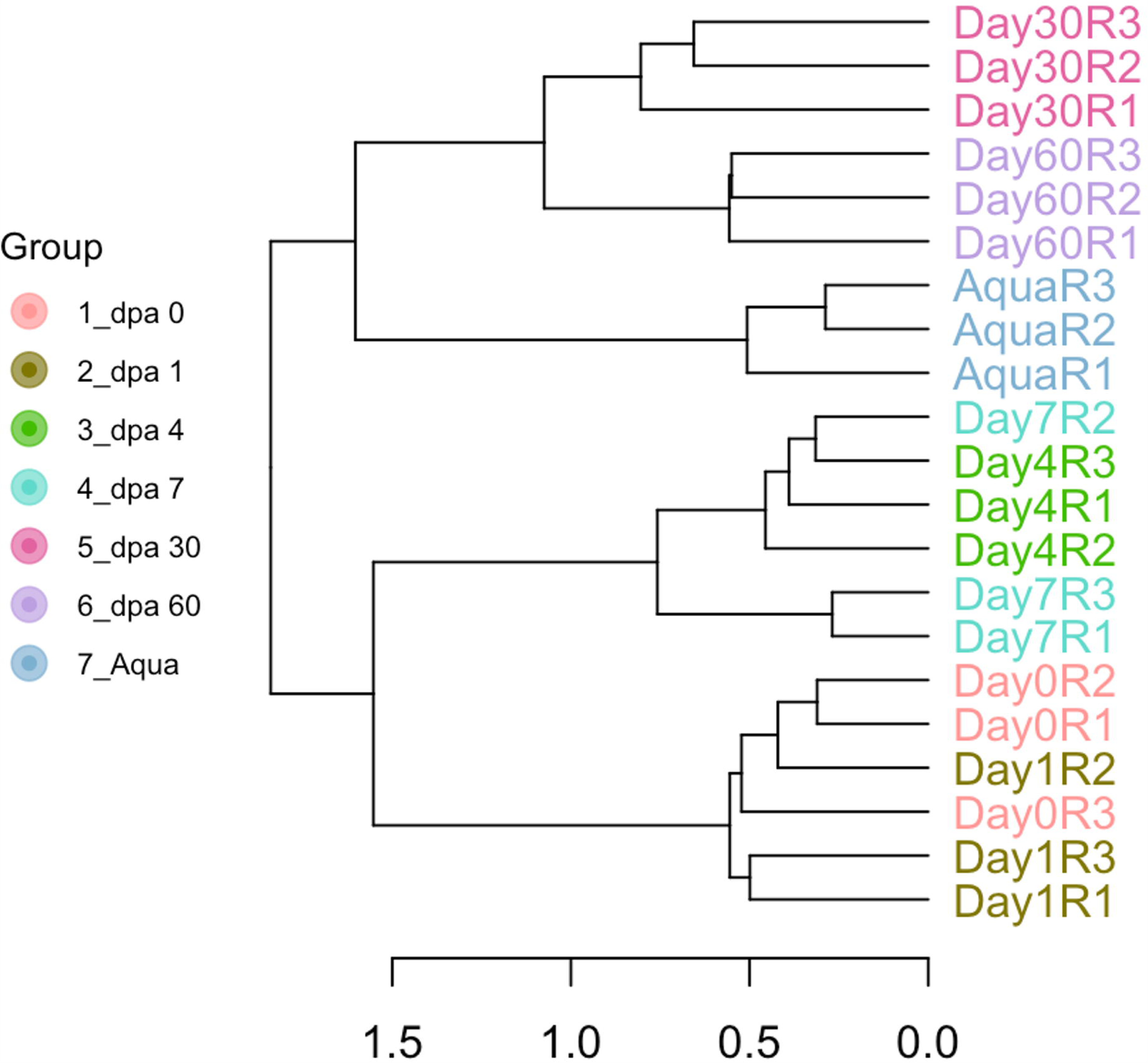

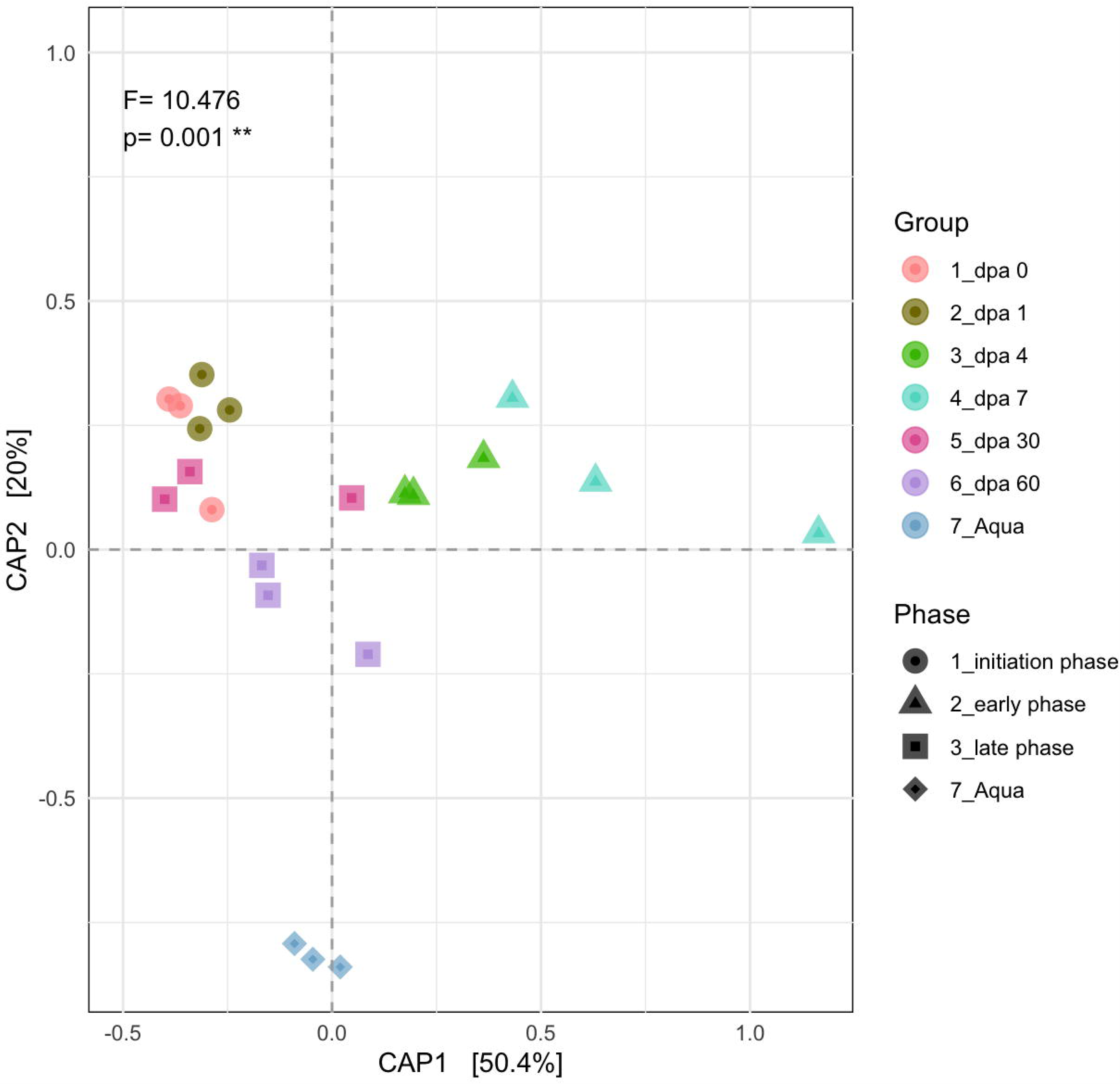

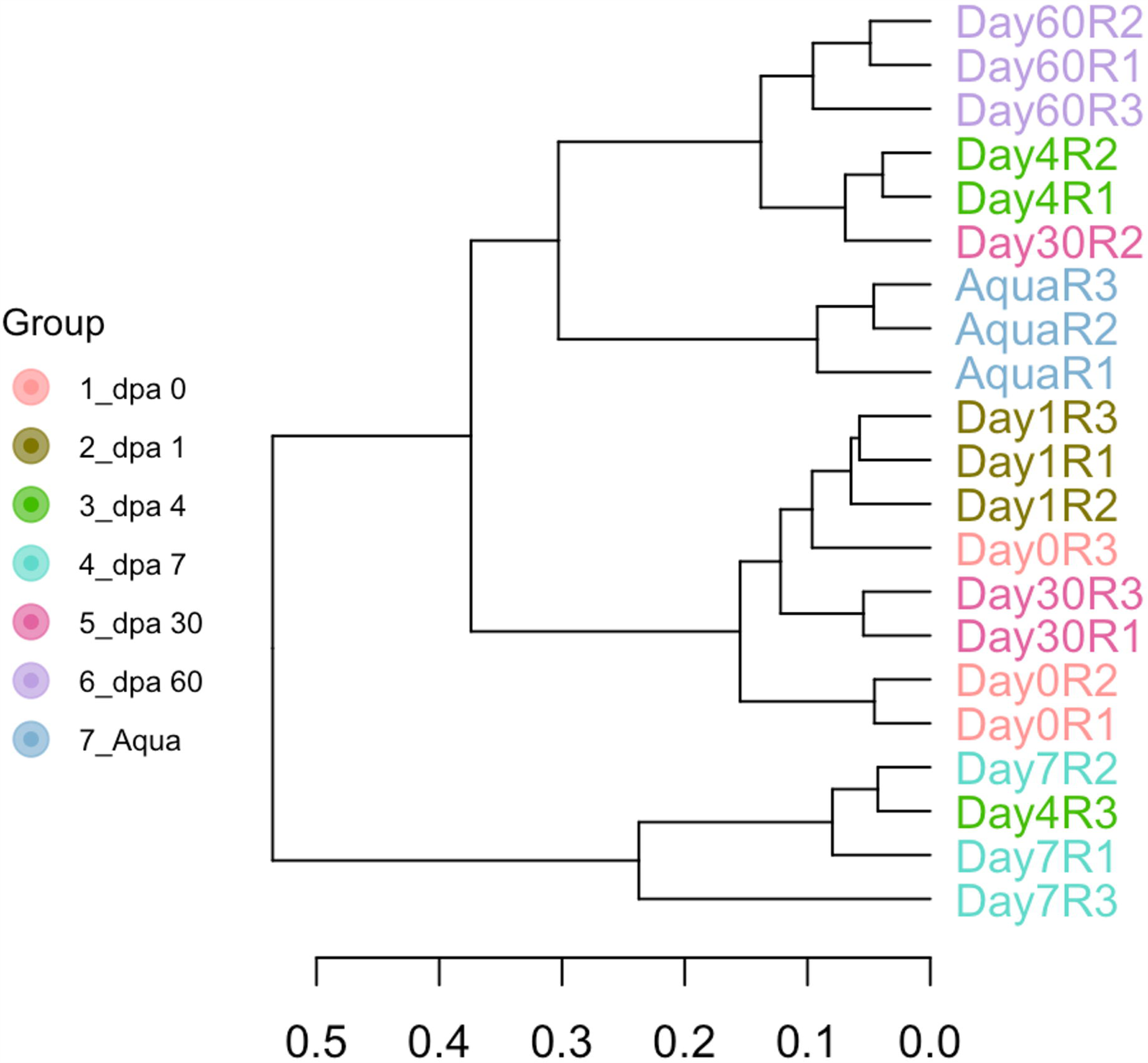

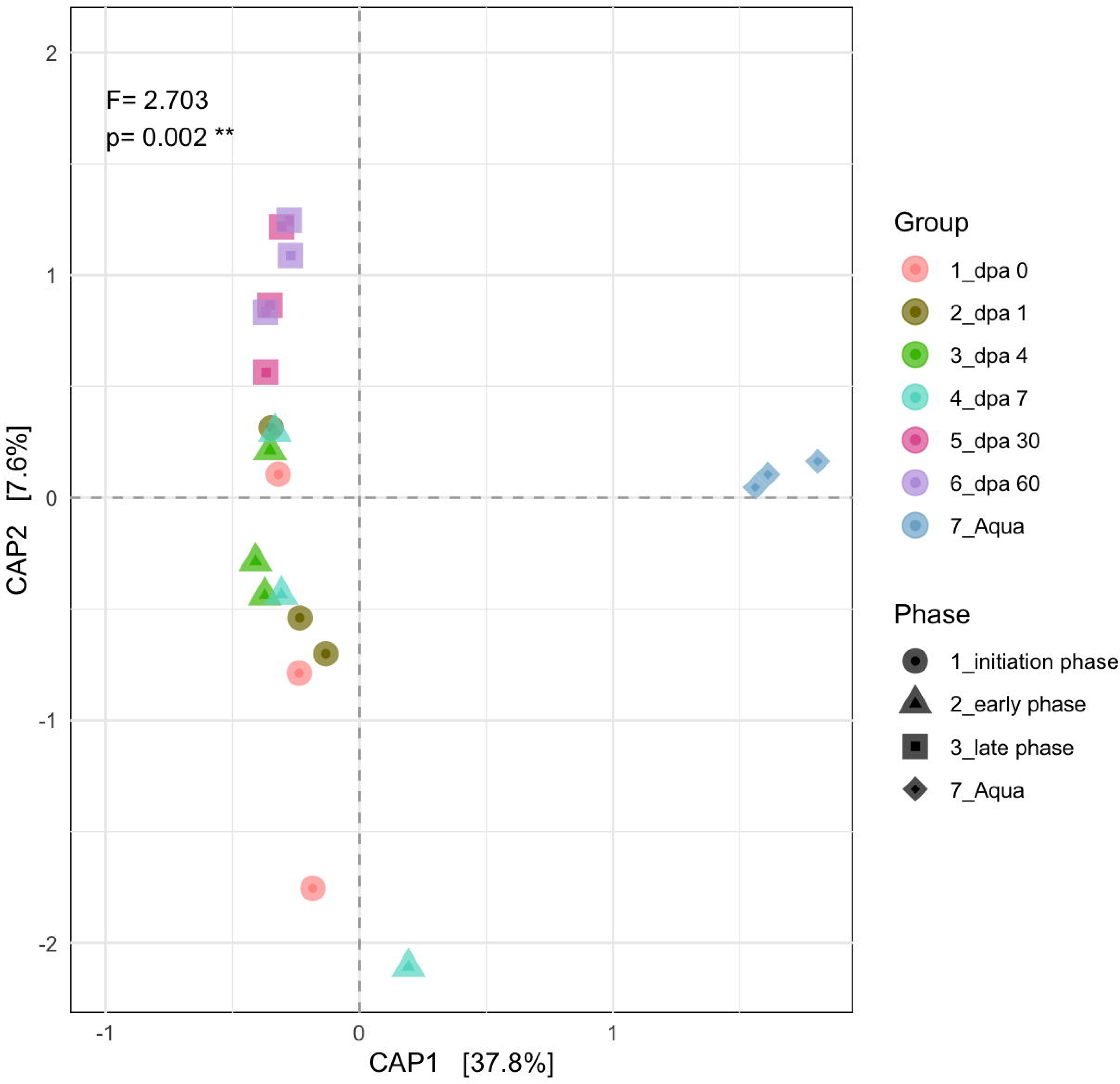

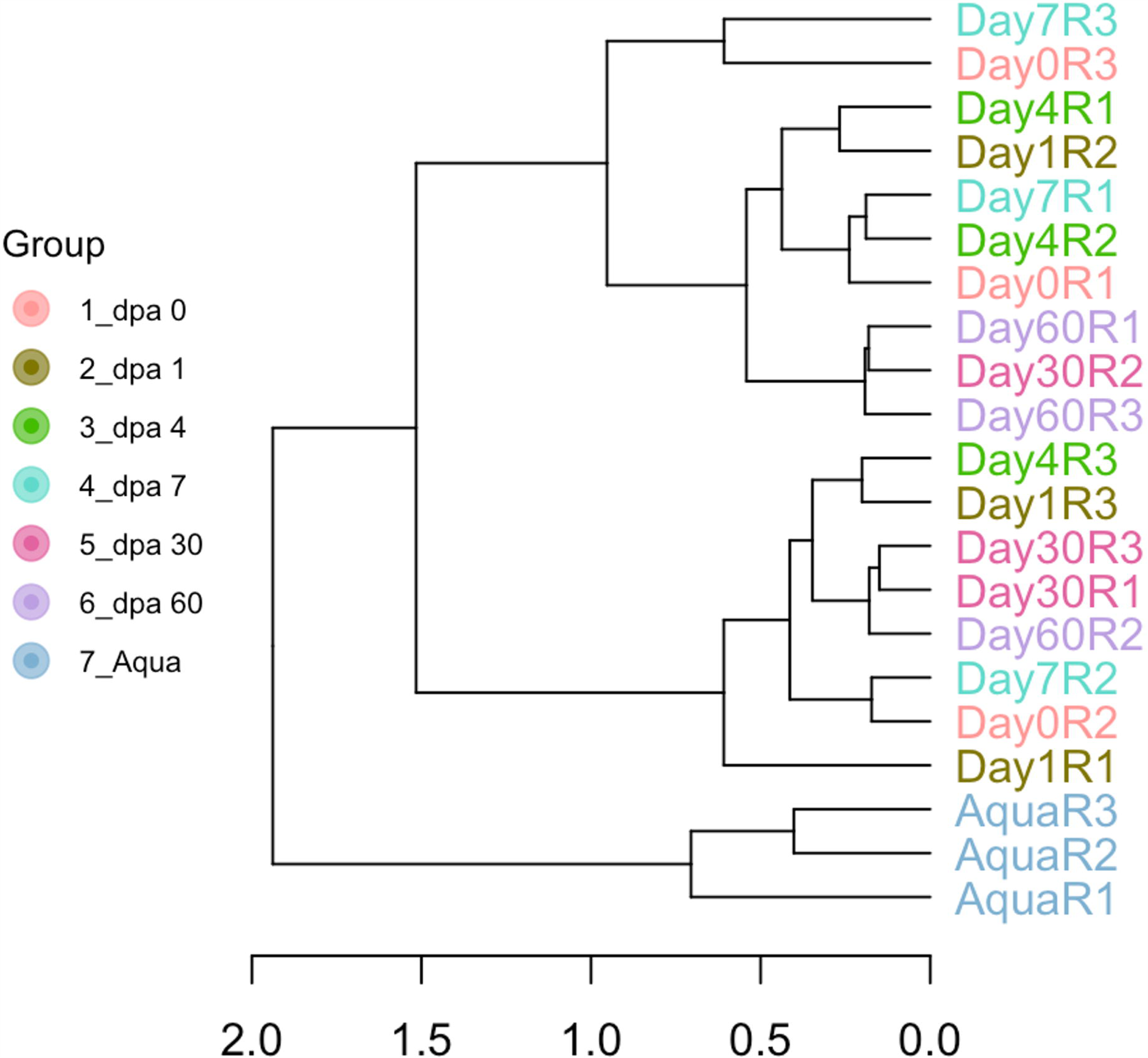

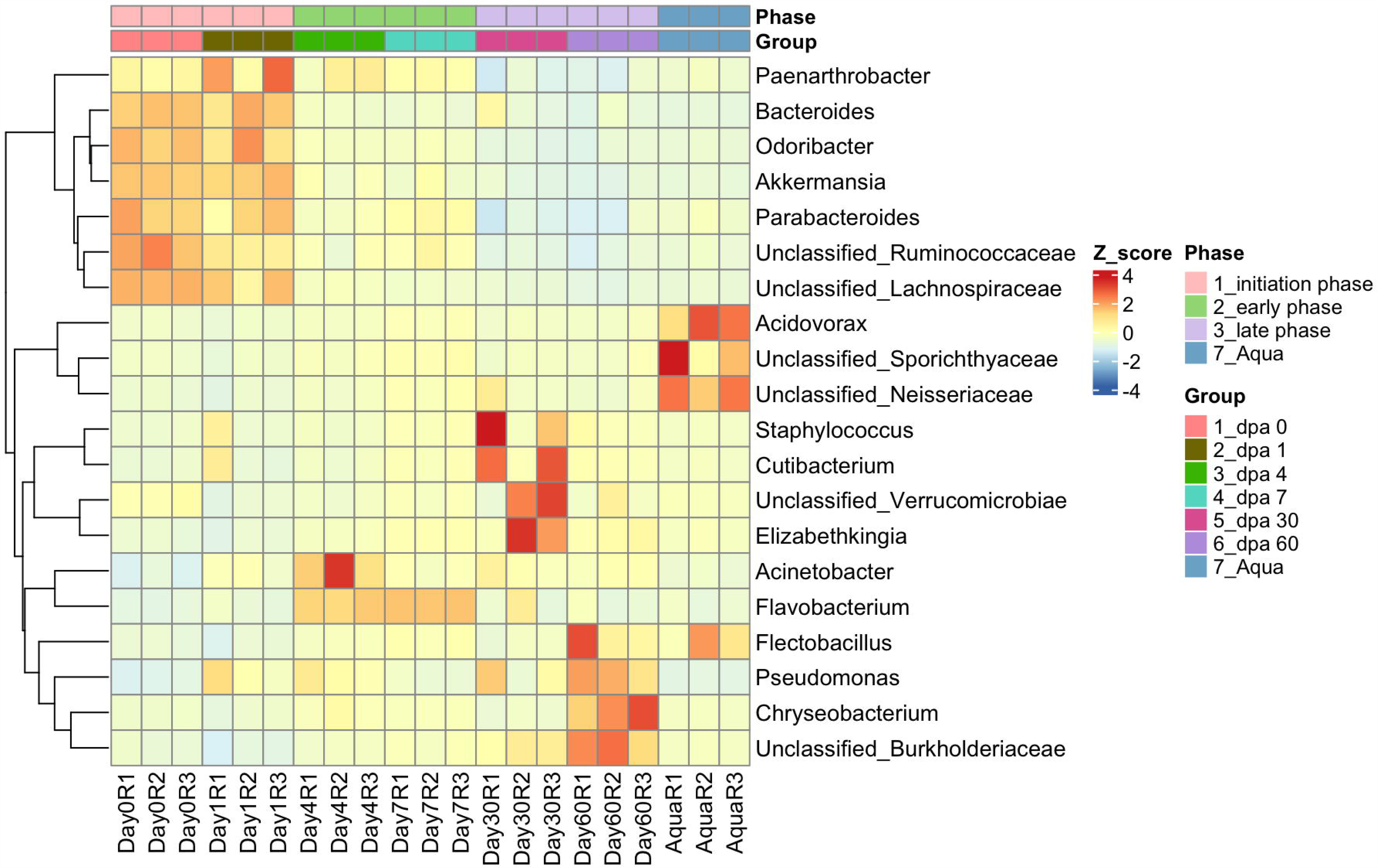

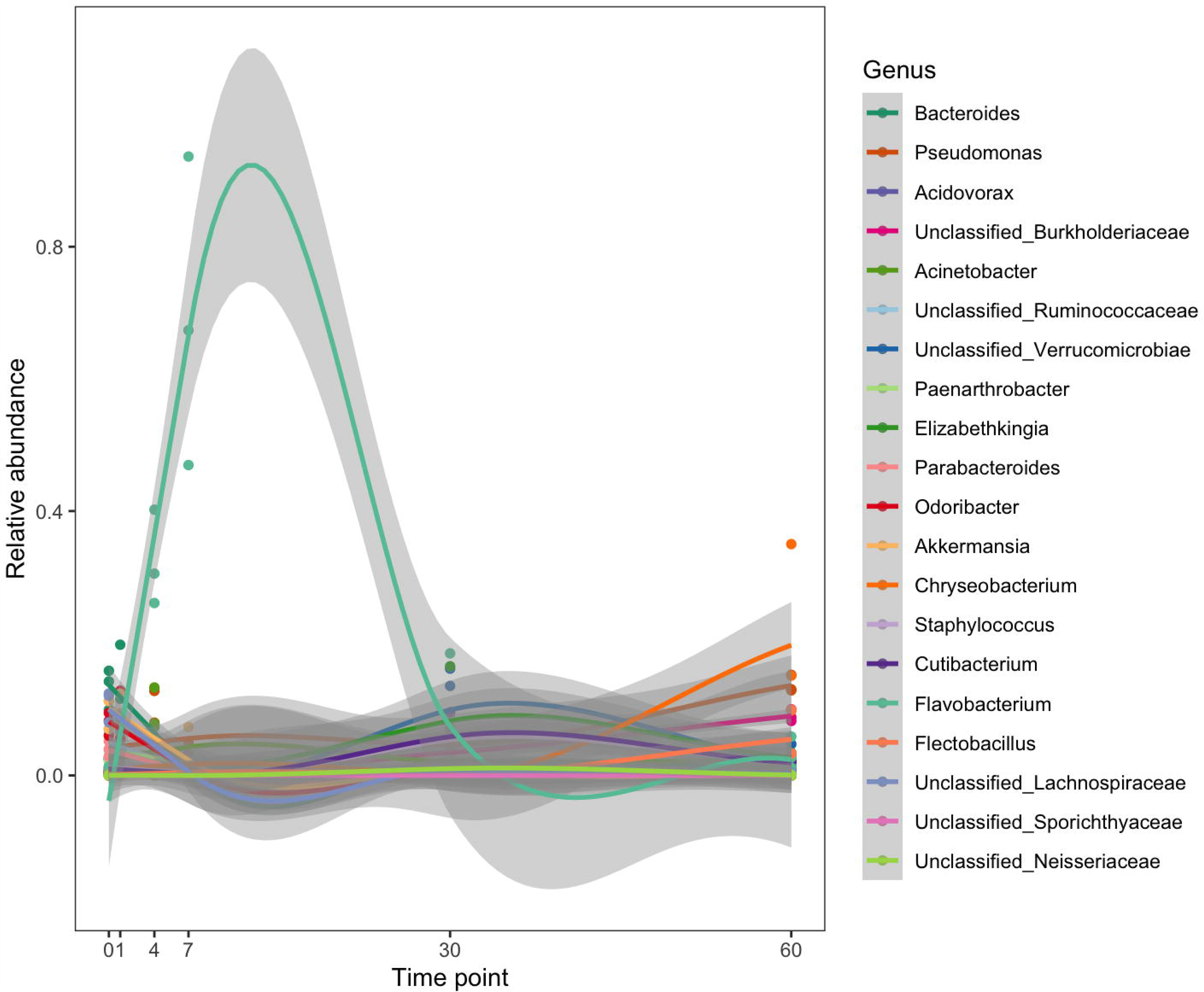

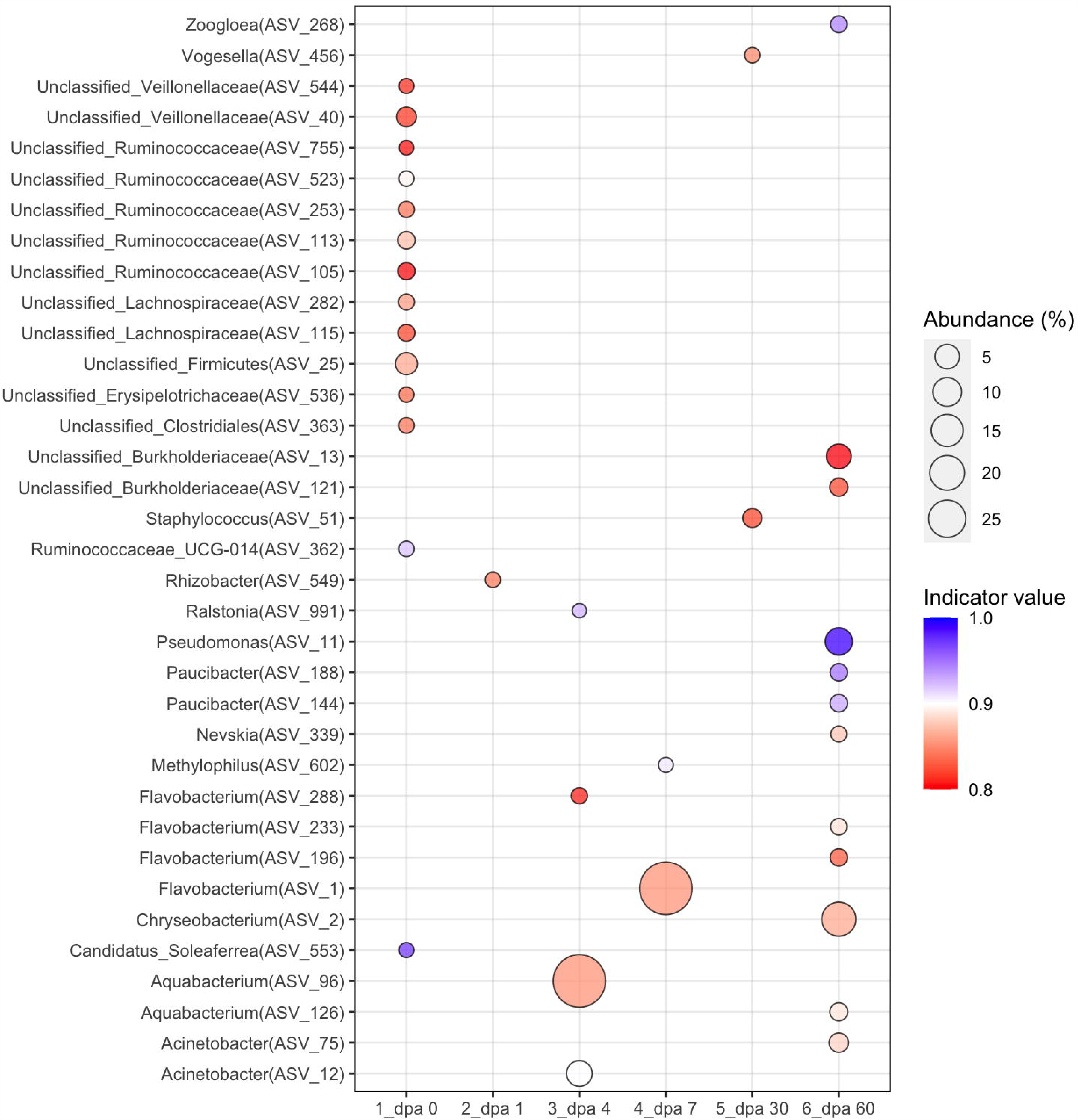

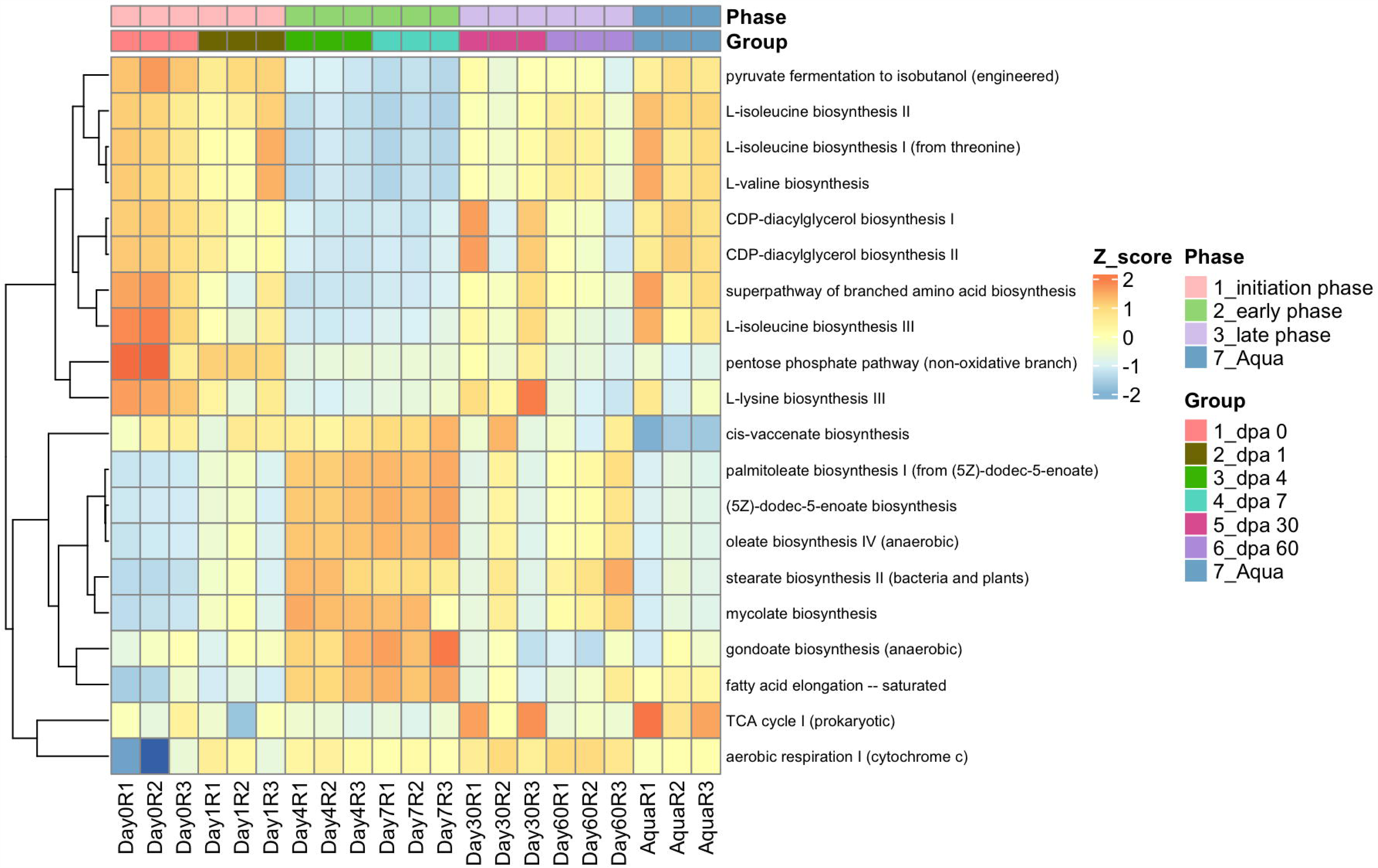

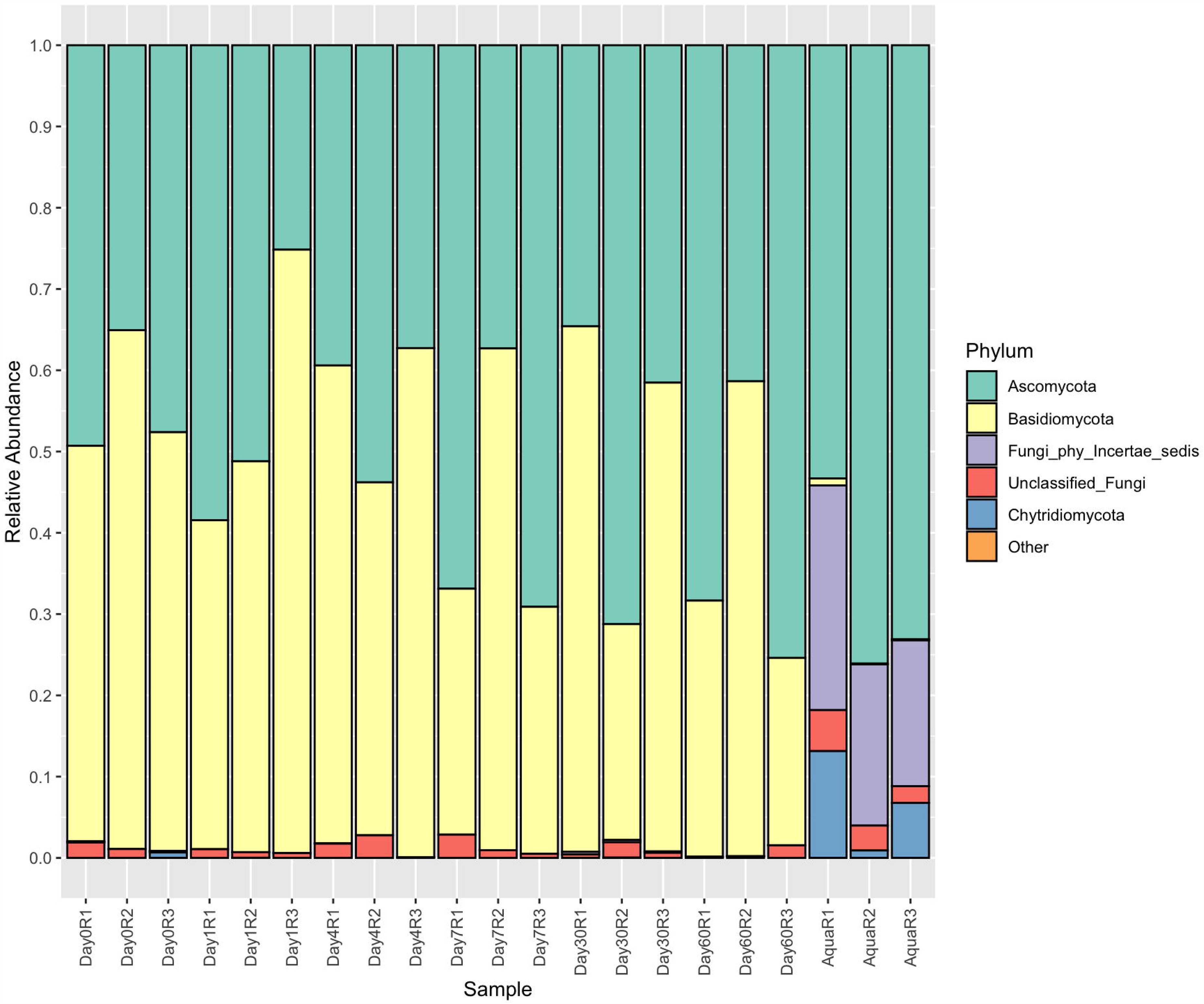

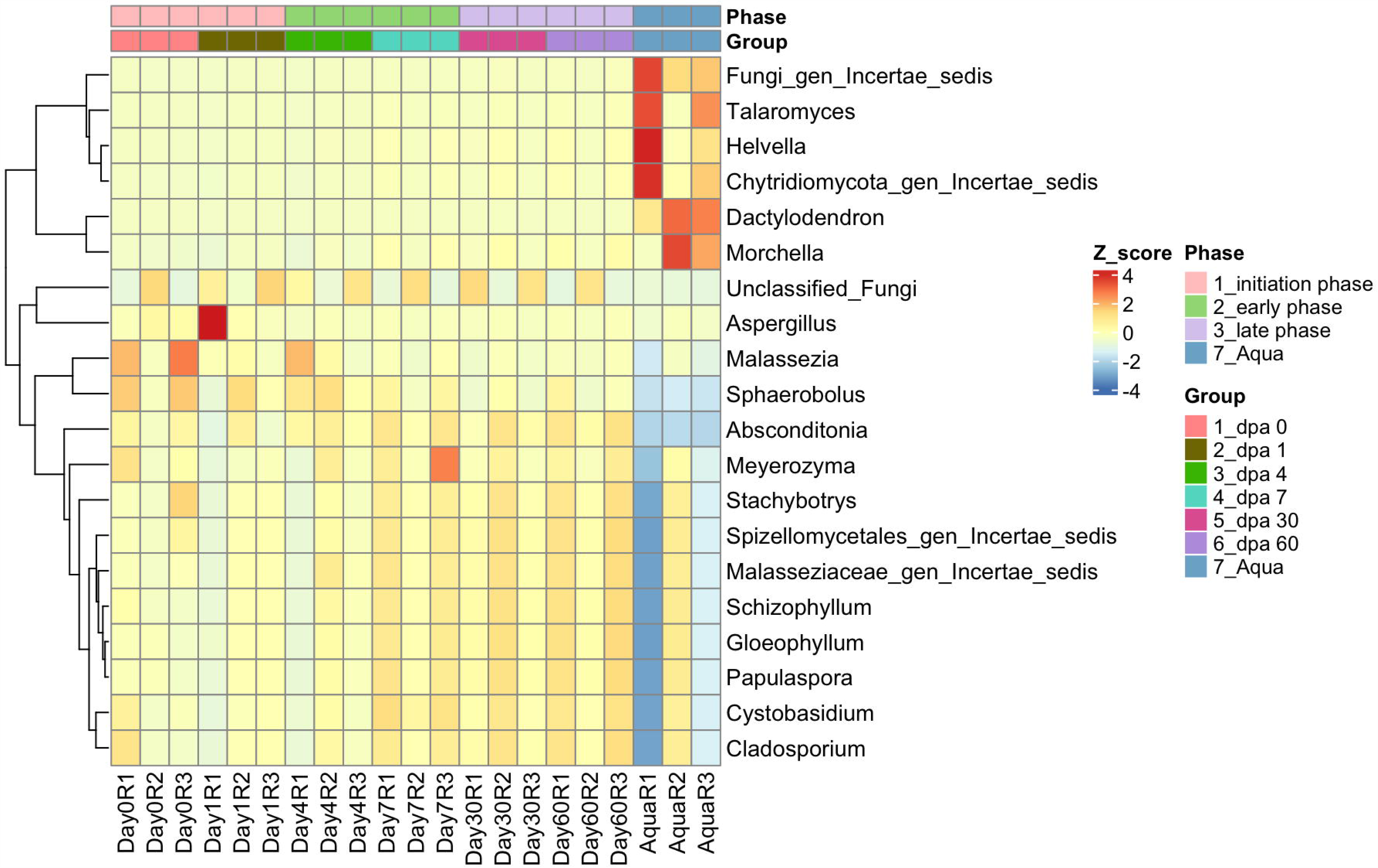

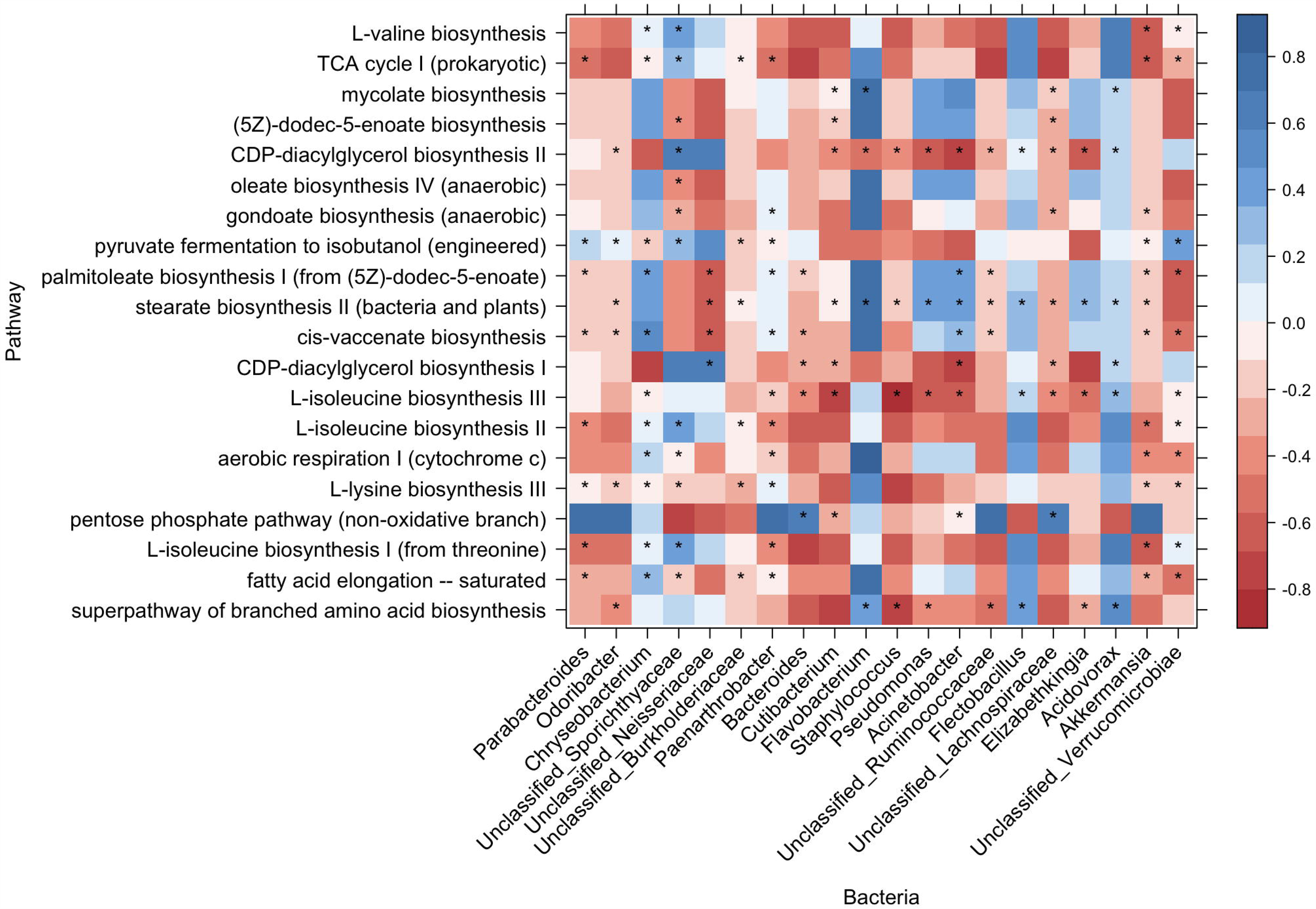

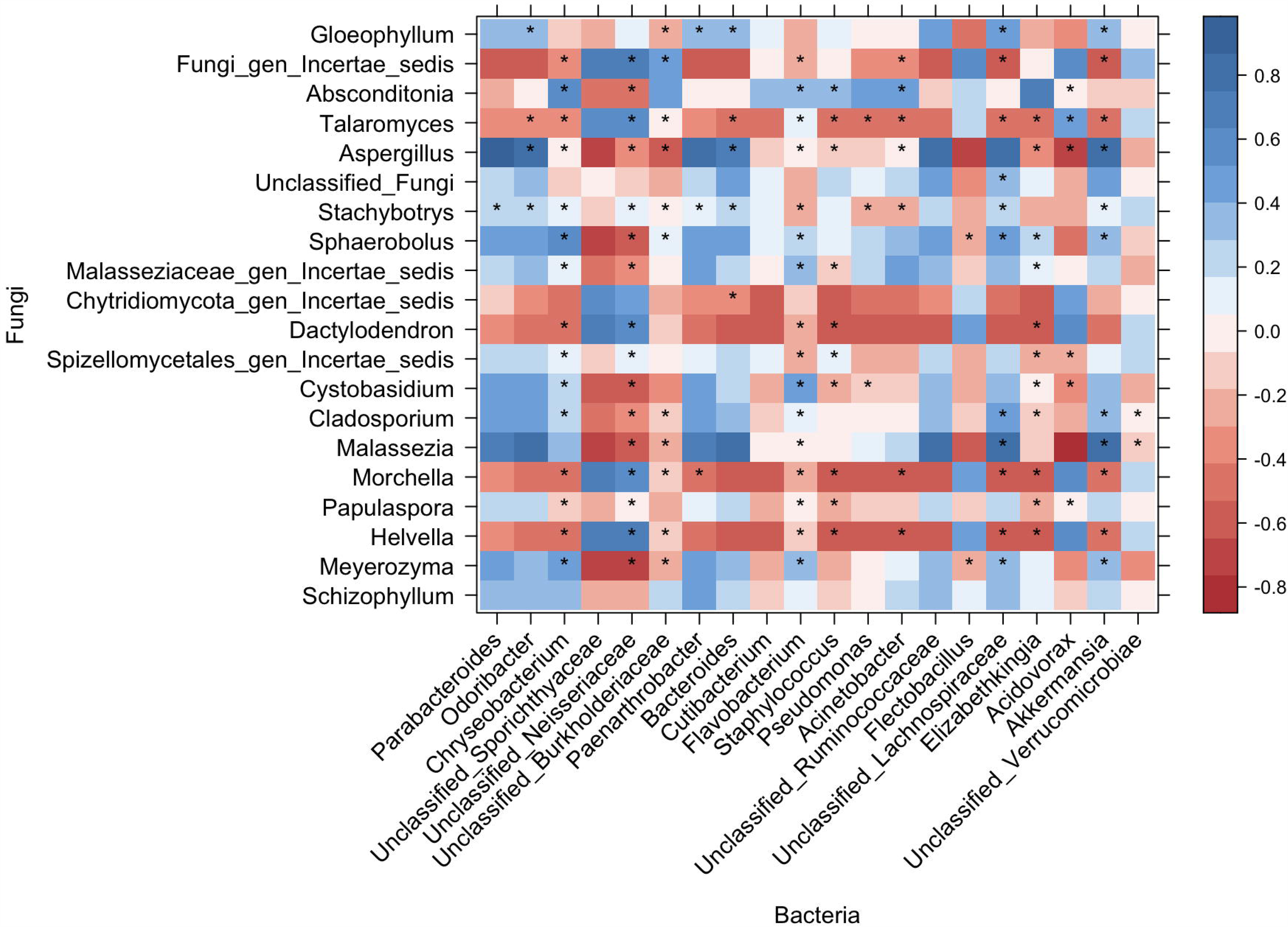

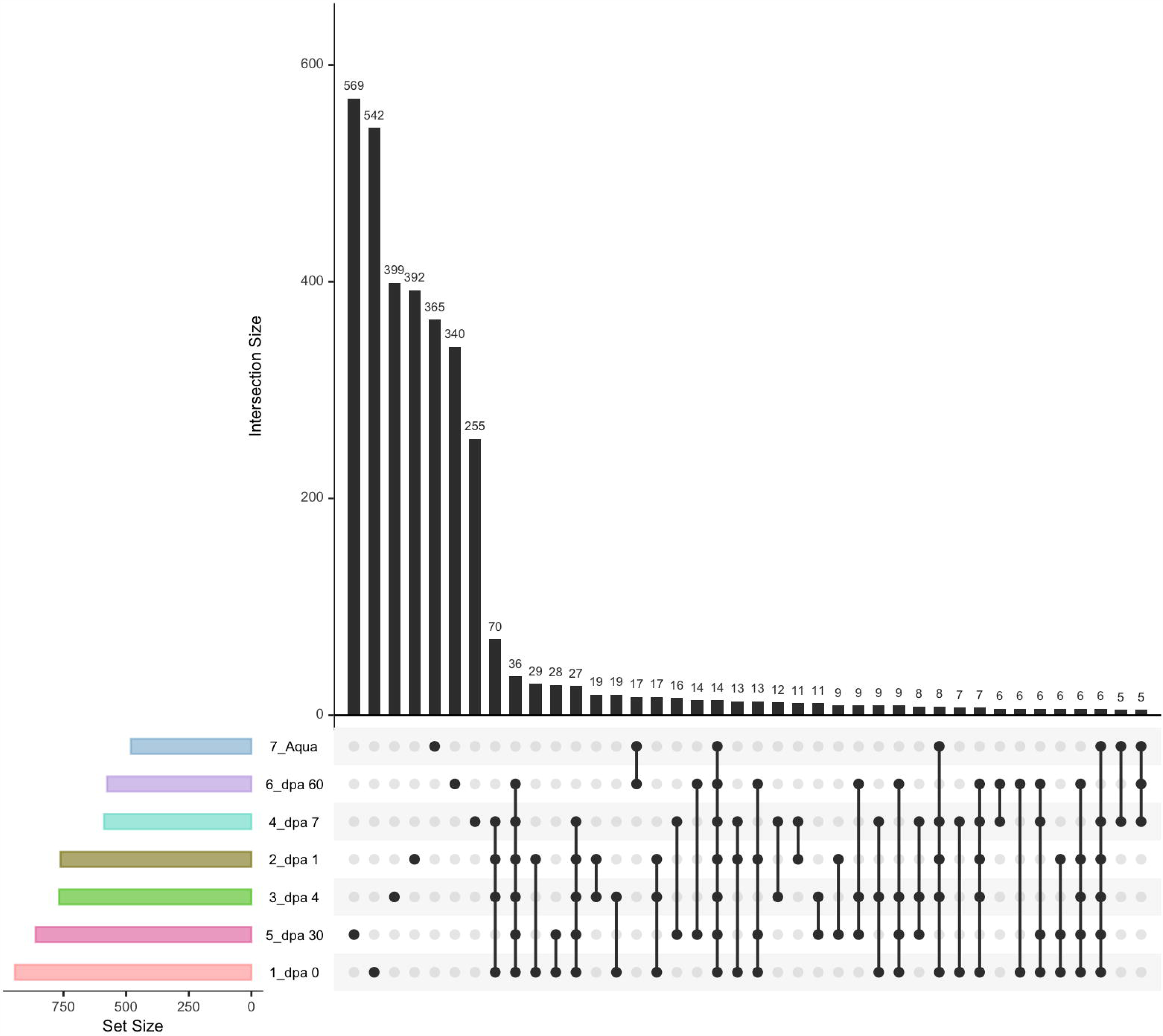

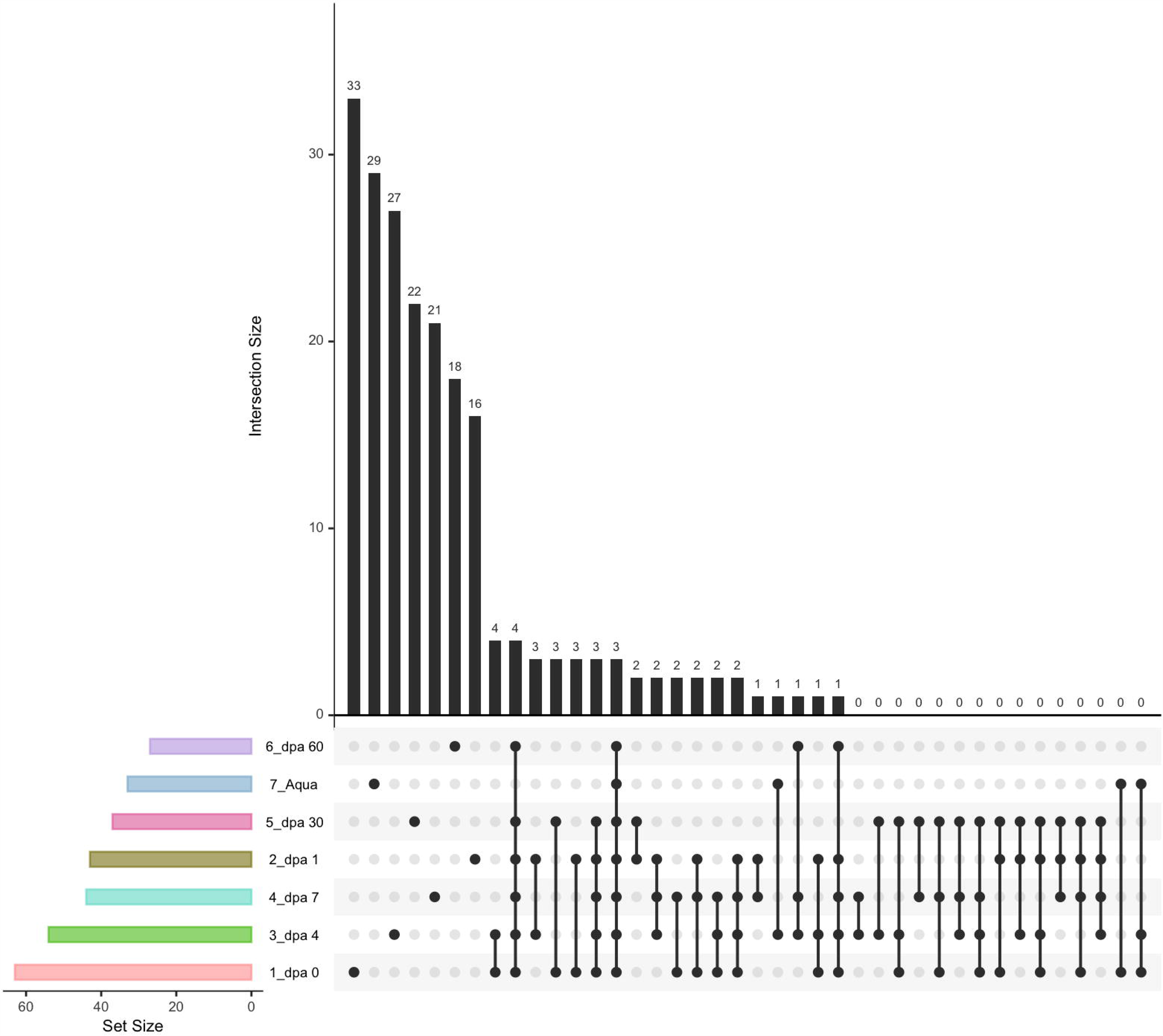

