## Supplemental Tables and Figures for "Temporal Microbiome Changes in Axolotl Limb Regeneration: Stage-Specific Restructuring of Bacterial and Fungal Communities with a *Flavobacterium* Bloom During Blastema Proliferation"

**Table S1.1. The observed, Chao1, Shannon, and Simpson metrics were utilized to assess the alpha diversity within the bacterial communities of each study group.** To determine the variations in alpha diversity between samples, the Kruskal-Wallis test was employed.  Numbers with * indicate statistically significant p values.

| **Measure Types** | **chi-squared** | **df** | **p-value** |
| --- | --- | --- | --- |
| **Observed** | 9.517 | 6 | 0.1465 |
| **Chao1** | 9.4892 | 6 | 0.1539 |
| **Shannon** | 16 | 6 | 0.01375^*^ |
| **Simpson** | 15.913 | 6 | 0.01423^*^ |

**p: 0 ‘***’ 0.001 ‘**’ 0.01 ‘*’ 0.05 ‘.’ 0.1 ‘ ’ 1**

**Table S1.2. The observed, Chao1, Shannon, and Simpson metrics were utilized to assess the alpha diversity within the fungal communities of each study group.** To determine the variations in alpha diversity between samples, the Kruskal-Wallis test was employed.  Numbers with * indicate statistically significant p values.

| **Measure Types** | **chi-squared** | **df** | **p-value** |
| --- | --- | --- | --- |
| **Observed** | 13.785 | 6 | 0.03213* |
| **Chao1** | 13.325 | 6 | 0.03816* |
| **Shannon** | 15.55 | 6 | 0.01639* |
| **Simpson** | 12.502 | 6 | 0.05166 |

**p: 0 ‘***’ 0.001 ‘**’ 0.01 ‘*’ 0.05 ‘.’ 0.1 ‘ ’ 1**

**Table S2.1. The observed, Chao1, Shannon, and Simpson metrics were utilized to assess the alpha diversity within the bacterial communities of each study group.** The post-hoc Dunn’s test were used to identify alpha diversity differences between samples. Numbers with * indicate statistically significant p values for each comparison.

| **Comparison of x by group (Observed)** | **p** | **Comparison of x by group (Chao1)** | **p** |
| --- | --- | --- | --- |
| 1_dpa 0-2_dpa 1 | 0.2553 | 1_dpa 0-2_dpa 1 | 0.2346 |
| 1_dpa 0-3_dpa 4 | 0.2993 | 1_dpa 0-3_dpa 4 | 0.2553 |
| 1_dpa 0-4_dpa 7 | 0.0500 | 1_dpa 0-4_dpa 7 | 0.0651* |
| 1_dpa 0-5_dpa 30 | 0.1785 | 1_dpa 0-5_dpa 30 | 0.1962 |
| 1_dpa 0-6_dpa 60 | 0.0106** | 1_dpa 0-6_dpa 60 | 0.0150* |
| 1_dpa 0-7_Aqua | 0.0126** | 1_dpa 0-7_Aqua | 0.0089** |
| 2_dpa 1-3_dpa 4 | 0.4477 | 2_dpa 1-3_dpa 4 | 0.4738 |
| 2_dpa 1-4_dpa 7 | 0.1618 | 2_dpa 1-4_dpa 7 | 0.2149 |
| 2_dpa 1-5_dpa 30 | 0.3962 | 2_dpa 1-5_dpa 30 | 0.4477 |
| 2_dpa 1-6_dpa 60 | 0.0500 | 2_dpa 1-6_dpa 60 | 0.0739 |
| 2_dpa 1-7_Aqua | 0.0572 | 2_dpa 1-7_Aqua | 0.0500 |
| 3_dpa 4-4_dpa 7 | 0.1317 | 3_dpa 4-4_dpa 7 | 0.1962 |
| 3_dpa 4-5_dpa 30 | 0.3465 | 3_dpa 4-5_dpa 30 | 0.4218 |
| 3_dpa 4-6_dpa 60 | 0.0378 | 3_dpa 4-6_dpa 60 | 0.0651 |
| 3_dpa 4-7_Aqua | 0.0436 | 3_dpa 4-7_Aqua | 0.0436 |
| 4_dpa 7-5_dpa 30 | 0.2346 | 4_dpa 7-5_dpa 30 | 0.2553 |
| 4_dpa 7-6_dpa 60 | 0.2553 | 4_dpa 7-6_dpa 60 | 0.2553 |
| 4_dpa 7-7_Aqua | 0.2769 | 4_dpa 7-7_Aqua | 0.1962 |
| 5_dpa 30-6_dpa 60 | 0.0835 | 5_dpa 30-6_dpa 60 | 0.0941 |
| 5_dpa 30-7_Aqua | 0.0941 | 5_dpa 30-7_Aqua | 0.0651 |
| 6_dpa 60-7_Aqua | 0.4738 | 6_dpa 60-7_Aqua | 0.4218 |
| **Comparison of x by group (Shannon)** | **p** | **Comparison of x by group (Simpson)** | **p** |
| 1_dpa 0-2_dpa 1 | 0.4738 | 1_dpa 0-2_dpa 1 | 0.4218 |
| 1_dpa 0-3_dpa 4 | 0.0572* | 1_dpa 0-3_dpa 4 | 0.0282* |
| 1_dpa 0-4_dpa 7 | 0.0023** | 1_dpa 0-4_dpa 7 | 0.0015** |
| 1_dpa 0-5_dpa 30 | 0.2553 | 1_dpa 0-5_dpa 30 | 0.3226 |
| 1_dpa 0-6_dpa 60 | 0.0835* | 1_dpa 0-6_dpa 60 | 0.1181 |
| 1_dpa 0-7_Aqua | 0.0035** | 1_dpa 0-7_Aqua | 0.0242* |
| 2_dpa 1-3_dpa 4 | 0.0651* | 2_dpa 1-3_dpa 4 | 0.0176 |
| 2_dpa 1-4_dpa 7 | 0.0029** | 2_dpa 1-4_dpa 7 | 0.0008*** |
| 2_dpa 1-5_dpa 30 | 0.2769 | 2_dpa 1-5_dpa 30 | 0.2553 |
| 2_dpa 1-6_dpa 60 | 0.0941* | 2_dpa 1-6_dpa 60 | 0.0835* |
| 2_dpa 1-7_Aqua | 0.0042** | 2_dpa 1-7_Aqua | 0.0150 |
| 3_dpa 4-4_dpa 7 | 0.1056* | 3_dpa 4-4_dpa 7 | 0.1462 |
| 3_dpa 4-5_dpa 30 | 0.1785 | 3_dpa 4-5_dpa 30 | 0.0739* |
| 3_dpa 4-6_dpa 60 | 0.4218 | 3_dpa 4-6_dpa 60 | 0.2346 |
| 3_dpa 4-7_Aqua | 0.1317 | 3_dpa 4-7_Aqua | 0.4738 |
| 4_dpa 7-5_dpa 30 | 0.0150** | 4_dpa 7-5_dpa 30 | 0.0062** |
| 4_dpa 7-6_dpa 60 | 0.0739* | 4_dpa 7-6_dpa 60 | 0.0378* |
| 4_dpa 7-7_Aqua | 0.4477 | 4_dpa 7-7_Aqua | 0.1618 |
| 5_dpa 30-6_dpa 60 | 0.2346 | 5_dpa 30-6_dpa 60 | 0.2346 |
| 5_dpa 30-7_Aqua | 0.0207** | 5_dpa 30-7_Aqua | 0.0651* |
| 6_dpa 60-7_Aqua | 0.0941* | 6_dpa 60-7_Aqua | 0.2149 |

**p: 0 ‘***’ 0.001 ‘**’ 0.01 ‘*’ 0.05 ‘.’ 0.1 ‘ ’ 1**

**Table S2.2. The observed, Chao1, Shannon, and Simpson metrics were utilized to assess the alpha diversity within the fungal communities of each study group.** The post-hoc Dunn’s test were used to identify alpha diversity differences between samples. Numbers with * indicate statistically significant p values for each comparison.

| **Comparison of x by group (Observed)** | **p** | **Comparison of x by group (Chao1)** | **p** |
| --- | --- | --- | --- |
| 1_dpa 0-2_dpa 1 | 0.1458 | 1_dpa 0-2_dpa 1 | 0.1537 |
| 1_dpa 0-3_dpa 4 | 0.3708 | 1_dpa 0-3_dpa 4 | 0.4738 |
| 1_dpa 0-4_dpa 7 | 0.1177 | 1_dpa 0-4_dpa 7 | 0.1116 |
| 1_dpa 0-5_dpa 30 | 0.0136* | 1_dpa 0-5_dpa 30 | 0.0609 |
| 1_dpa 0-6_dpa 60 | 0.0017** | 1_dpa 0-6_dpa 60 | 0.0026** |
| 1_dpa 0-7_Aqua | 0.0136* | 1_dpa 0-7_Aqua | 0.0176* |
| 2_dpa 1-3_dpa 4 | 0.2342 | 2_dpa 1-3_dpa 4 | 0.1699 |
| 2_dpa 1-4_dpa 7 | 0.4476 | 2_dpa 1-4_dpa 7 | 0.4217 |
| 2_dpa 1-5_dpa 30 | 0.1243 | 2_dpa 1-5_dpa 30 | 0.2992 |
| 2_dpa 1-6_dpa 60 | 0.0301* | 2_dpa 1-6_dpa 60 | 0.0377* |
| 2_dpa 1-7_Aqua | 0.1243 | 2_dpa 1-7_Aqua | 0.1387 |
| 3_dpa 4-4_dpa 7 | 0.1957 | 3_dpa 4-4_dpa 7 | 0.1246 |
| 3_dpa 4-5_dpa 30 | 0.0301* | 3_dpa 4-5_dpa 30 | 0.0693 |
| 3_dpa 4-6_dpa 60 | 0.0046** | 3_dpa 4-6_dpa 60 | 0.0031** |
| 3_dpa 4-7_Aqua | 0.0301* | 3_dpa 4-7_Aqua | 0.0206* |
| 4_dpa 7-5_dpa 30 | 0.1534 | 4_dpa 7-5_dpa 30 | 0.3710 |
| 4_dpa 7-6_dpa 60 | 0.0403* | 4_dpa 7-6_dpa 60 | 0.0570 |
| 4_dpa 7-7_Aqua | 0.1534 | 4_dpa 7-7_Aqua | 0.1871 |
| 5_dpa 30-6_dpa 60 | 0.2342 | 5_dpa 30-6_dpa 60 | 0.1055 |
| 5_dpa 30-7_Aqua | 0.5000 | 5_dpa 30-7_Aqua | 0.2879 |
| 6_dpa 60-7_Aqua | 0.2342 | 6_dpa 60-7_Aqua | 0.2447 |
| **Comparison of x by group (Shannon)** | **p** | **Comparison of x by group (Simpson)** | **p** |
| 1_dpa 0-2_dpa 1 | 0.0739 | 1_dpa 0-2_dpa 1 | 0.1618 |
| 1_dpa 0-3_dpa 4 | 0.1962 | 1_dpa 0-3_dpa 4 | 0.3465 |
| 1_dpa 0-4_dpa 7 | 0.0500* | 1_dpa 0-4_dpa 7 | 0.0500* |
| 1_dpa 0-5_dpa 30 | 0.0042** | 1_dpa 0-5_dpa 30 | 0.0106* |
| 1_dpa 0-6_dpa 60 | 0.0005*** | 1_dpa 0-6_dpa 60 | 0.0029** |
| 1_dpa 0-7_Aqua | 0.2346 | 1_dpa 0-7_Aqua | 0.2553 |
| 2_dpa 1-3_dpa 4 | 0.2769 | 2_dpa 1-3_dpa 4 | 0.2769 |
| 2_dpa 1-4_dpa 7 | 0.4218 | 2_dpa 1-4_dpa 7 | 0.2553 |
| 2_dpa 1-5_dpa 30 | 0.1181 | 2_dpa 1-5_dpa 30 | 0.0941 |
| 2_dpa 1-6_dpa 60 | 0.0327* | 2_dpa 1-6_dpa 60 | 0.0378* |
| 2_dpa 1-7_Aqua | 0.2346 | 2_dpa 1-7_Aqua | 0.3711 |
| 3_dpa 4-4_dpa 7 | 0.2149 | 3_dpa 4-4_dpa 7 | 0.1056 |
| 3_dpa 4-5_dpa 30 | 0.0378* | 3_dpa 4-5_dpa 30 | 0.0282* |
| 3_dpa 4-6_dpa 60 | 0.0075** | 3_dpa 4-6_dpa 60 | 0.0089** |
| 3_dpa 4-7_Aqua | 0.4477 | 3_dpa 4-7_Aqua | 0.3962 |
| 4_dpa 7-5_dpa 30 | 0.1618 | 4_dpa 7-5_dpa 30 | 0.2553 |
| 4_dpa 7-6_dpa 60 | 0.0500* | 4_dpa 7-6_dpa 60 | 0.1317 |
| 4_dpa 7-7_Aqua | 0.1785 | 4_dpa 7-7_Aqua | 0.1618 |
| 5_dpa 30-6_dpa 60 | 0.2553 | 5_dpa 30-6_dpa 60 | 0.3226 |
| 5_dpa 30-7_Aqua | 0.0282* | 5_dpa 30-7_Aqua | 0.0500* |
| 6_dpa 60-7_Aqua | 0.0051** | 6_dpa 60-7_Aqua | 0.0176* |

**p: 0 ‘***’ 0.001 ‘**’ 0.01 ‘*’ 0.05 ‘.’ 0.1 ‘ ’ 1**

**Table S3. Pairwise PERMANOVA comparisons between sample types in bacterial communities.** Numbers with * indicate statistically significant p values for each comparison. (Number of permutations: 999)

| **Comparison of x by group** | **t** | **P(perm)** |
| --- | --- | --- |
| 1_dpa 0-2_dpa 1 | 1,248 | 0,099 |
| 1_dpa 0-3_dpa 4 | 2,1906 | 0,094 |
| 1_dpa 0-4_dpa 7 | 2,1306 | 0,087 |
| 1_dpa 0-5_dpa 30 | 2,0136 | 0,091 |
| 1_dpa 0-6_dpa 60 | 2,5509 | 0,083 |
| 1_dpa 0-7_Aqua | 4,3173 | 0,098 |
| 2_dpa 1-3_dpa 4 | 1,7468 | 0,105 |
| 2_dpa 1-4_dpa 7 | 1,852 | 0,09 |
| 2_dpa 1-5_dpa 30 | 1,8894 | 0,102 |
| 2_dpa 1-6_dpa 60 | 2,3364 | 0,112 |
| 2_dpa 1-7_Aqua | 3,9512 | 0,097 |
| 3_dpa 4-4_dpa 7 | 1,3798 | 0,099 |
| 3_dpa 4-5_dpa 30 | 1,881 | 0,119 |
| 3_dpa 4-6_dpa 60 | 2,291 | 0,112 |
| 3_dpa 4-7_Aqua | 4,231 | 0,106 |
| 4_dpa 7-5_dpa 30 | 1,6932 | 0,108 |
| 4_dpa 7-6_dpa 60 | 2,0455 | 0,101 |
| 4_dpa 7-7_Aqua | 3,1939 | 0,084 |
| 5_dpa 30-6_dpa 60 | 1,4176 | 0,086 |
| 5_dpa 30-7_Aqua | 2,5746 | 0,108 |
| 6_dpa 60-7_Aqua | 2,9739 | 0,088 |

**t, t-statistic; P, probability; Perms, number of unique permutations; significant results are numbers with * indicate.**

**Table S4. Pairwise PERMANOVA comparisons between sample types in fungal communities.** Numbers with * indicate statistically significant p values for each comparison. (Number of permutations: 999)

| **Comparison of x by group** | **t** | **P(perm)** |
| --- | --- | --- |
| 1_dpa 0-2_dpa 1 | 1,0737 | 0,3986 |
| 1_dpa 0-3_dpa 4 | 1,7773 | 0,102 |
| 1_dpa 0-4_dpa 7 | 0,97851 | 0,5012 |
| 1_dpa 0-5_dpa 30 | 1,5405 | 0,1012 |
| 1_dpa 0-6_dpa 60 | 0,90453 | 0,4929 |
| 1_dpa 0-7_Aqua | 1,4419 | 0,1983 |
| 2_dpa 1-3_dpa 4 | 1,0157 | 0,399 |
| 2_dpa 1-4_dpa 7 | 1,4716 | 0,1963 |
| 2_dpa 1-5_dpa 30 | 1,2336 | 0,3077 |
| 2_dpa 1-6_dpa 60 | 1,475 | 0,1977 |
| 2_dpa 1-7_Aqua | 1,4596 | 0,2032 |
| 3_dpa 4-4_dpa 7 | 1,7025 | 0,2062 |
| 3_dpa 4-5_dpa 30 | 1,369 | 0,1992 |
| 3_dpa 4-6_dpa 60 | 0,80908 | 0,8016 |
| 3_dpa 4-7_Aqua | 1,2495 | 0,2947 |
| 4_dpa 7-5_dpa 30 | 1,0737 | 0,3986 |
| 4_dpa 7-6_dpa 60 | 1,7773 | 0,102 |
| 4_dpa 7-7_Aqua | 0,97851 | 0,5012 |
| 5_dpa 30-6_dpa 60 | 1,5405 | 0,1012 |
| 5_dpa 30-7_Aqua | 0,90453 | 0,4929 |
| 6_dpa 60-7_Aqua | 1,4419 | 0,1983 |

**t, t-statistic; P, probability; Perms, number of unique permutations; significant results are numbers with * indicate.**

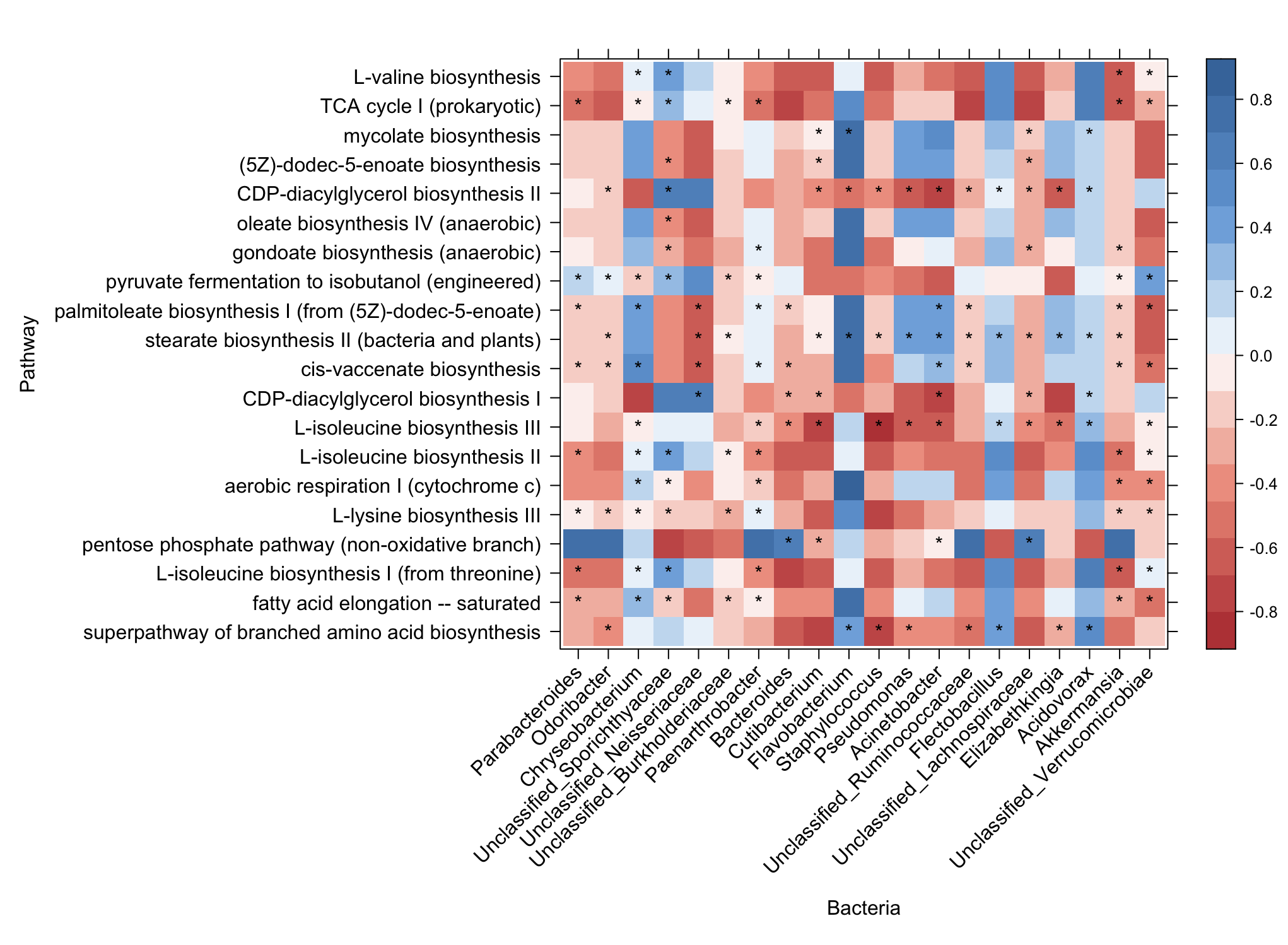

**Figure S1. Heatmap depicting Spearman Correlation Analysis Between Bacterial Taxa and Predicted Pathway Abundances.** Correlation coefficients are color-coded from dark red (indicating negative correlation) to dark blue (indicating positive correlation). Statistically significant correlations (|r| ≥ 0.40, p < 0.05) between bacterial taxa and pathway abundances are denoted by single asterisks (*).

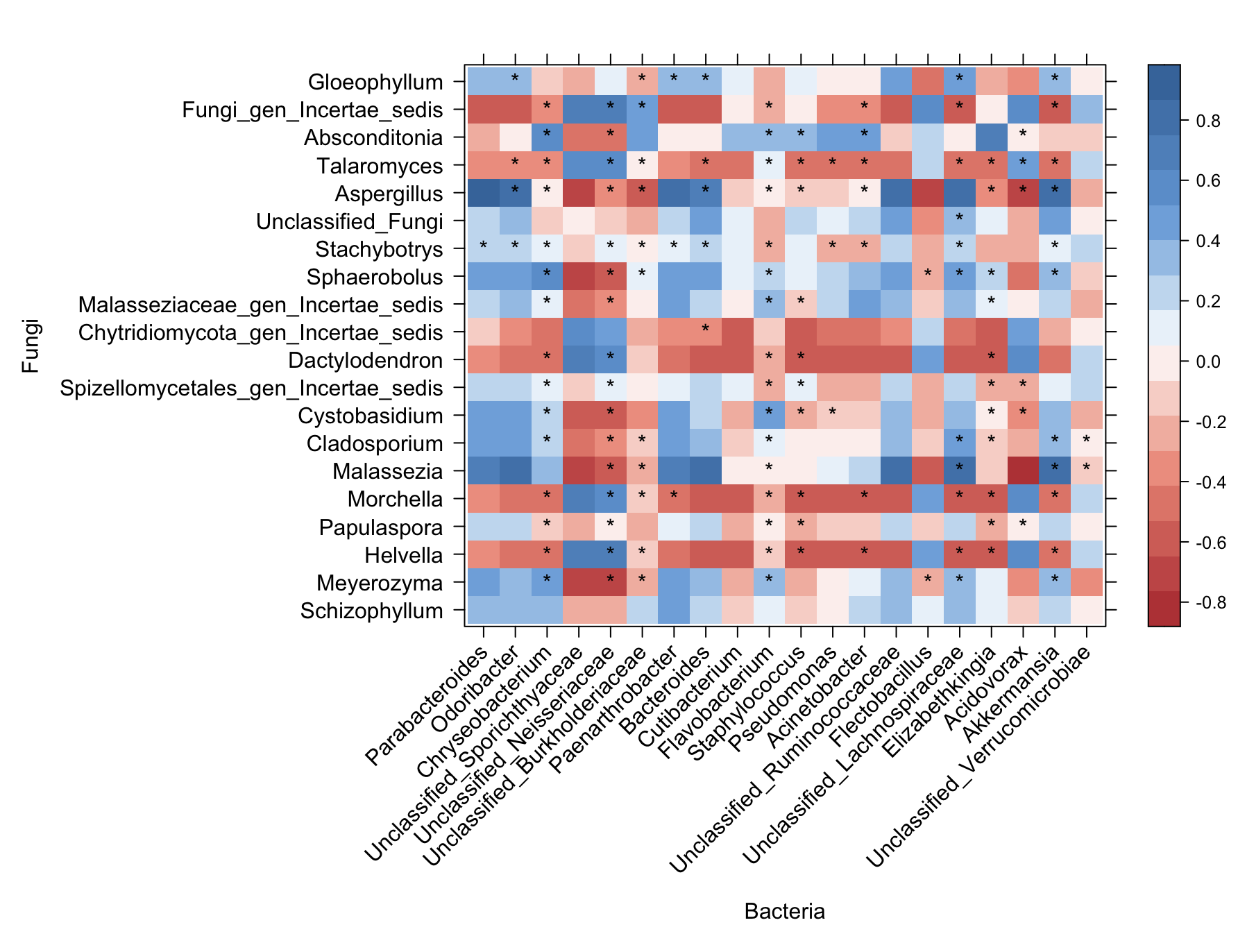

**Figure S2. Spearman Correlation analysis heatmap of bacterial and fungal Top20 genera.** Correlation coefficients are colored from dark red (negative correlation) to dark blue (positive correlation). Statistically significant (|r| ≥ 0.40, p < 0.05) correlations between bacterial and fungal taxa are shown as single (*).

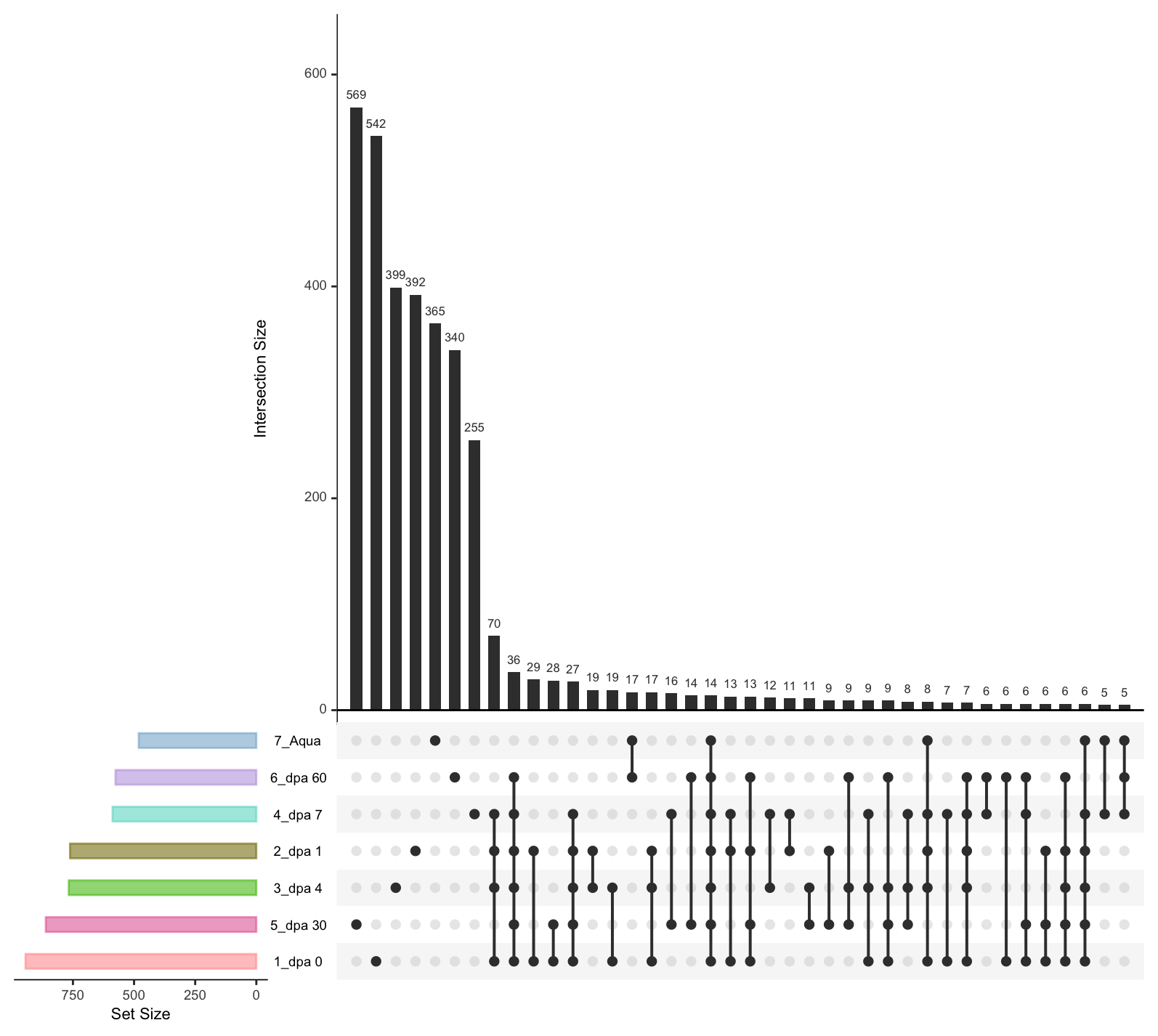

Figure S3A. **Number of Shared and Unique ASVs Between Longitudinally Obtained Samples.** (A) Upset plot illustrating shared and unique bacterial ASVs between samples.

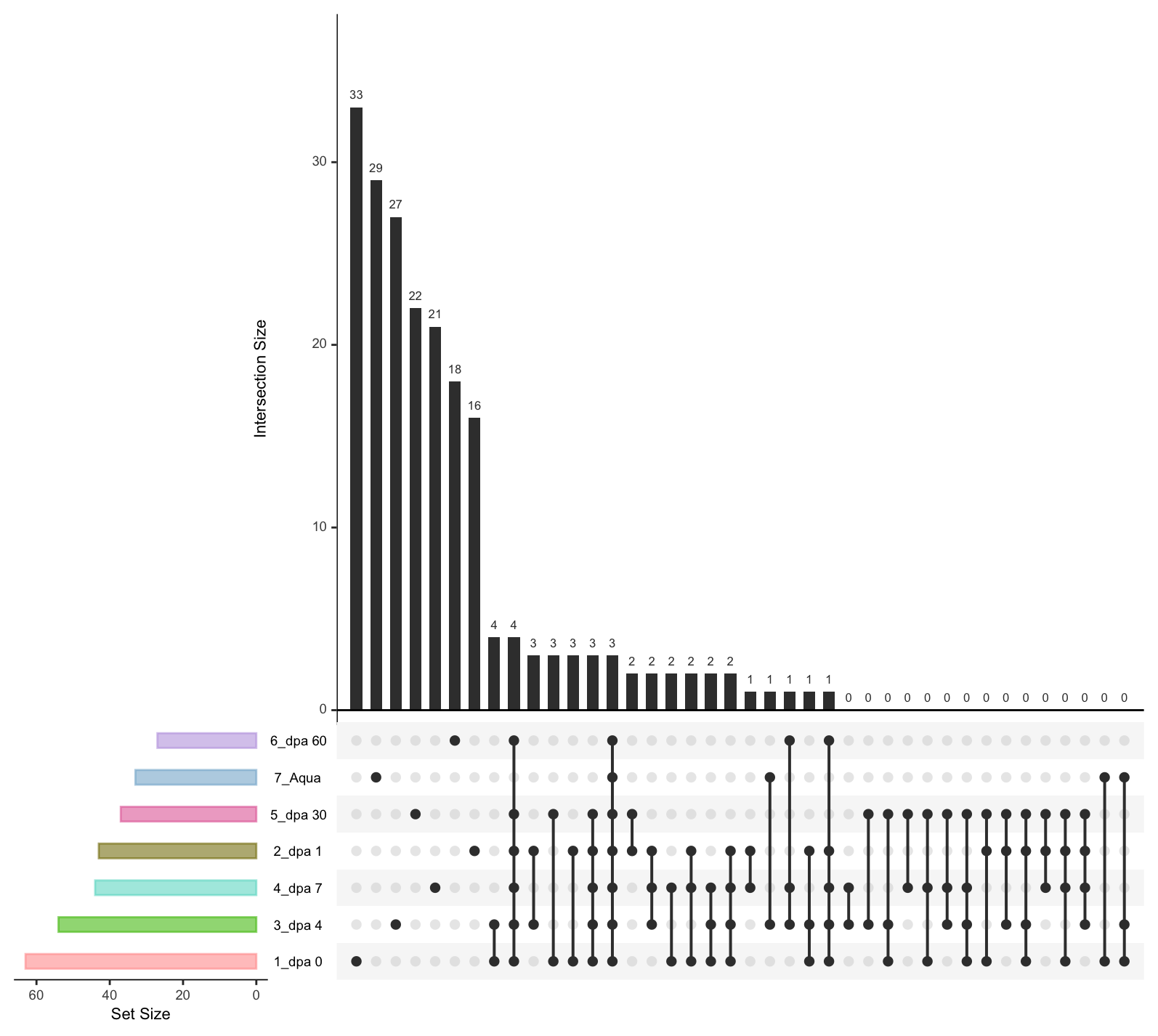

Figure S3B. **Number of Shared and Unique ASVs Between Longitudinally Obtained Samples.** Upset plot depicting shared and unique fungal ASVs. The size of each study group is represented in the left barplot. The overlapping black lines indicate the number of ASVs for each study group.
